# ImpuT2T: Pangenome-Based Patching for Human Genome Assemblies

**DOI:** 10.64898/2026.07.27.741037

**Authors:** Mao-Jan Lin, Vikram S. Shivakumar, Ben Langmead, Human Pangenome Reference Consortium

## Abstract

With improvements in sequencing and assembly have come many high-quality telomere-to-telomere assemblies and reference pangenomes. However, the long-read sequencing recipes needed for high quality assemblies are expensive, and out of reach for many research groups. Here we propose ImpuT2T, a method that takes an assembly produced via inexpensive HiFi sequencing reads, and uses a panel of T2T (or near-T2T) assemblies to scaffold and fill (“patch”) the gaps between the HiFi contigs. Benchmarking against reference assemblies demonstrates that ImpuT2T is highly effective at patching human HiFi assemblies, consistently outperforming existing patching approaches. Moreover, we show that including more haplotypes in the pangenome improves the quality of the patched assemblies, with the greatest gains achieved using the full HPRC Release 2 pangenome.

## 1 Background

Personal genome assembly offers significant advantages over using a standard reference genome, primarily by reducing reference bias that arises when the reference differs from the donor genome [1, 2]. However, generating high-quality personal assemblies remains challenging. Standard ∽ 30× long read whole-genome sequencing (WGS) followed by *de novo* assemblers such as hifiasm often results in relatively fragmented assemblies containing hundreds or thousands of contigs [3].

Since the assembly of the first complete telomere-to-telomere (T2T) human genome [4], a growing number of high-quality T2T or “near-T2T” human genomes have been produced [5, 6, 7, 8, 9]. Further, large-scale efforts such as the Human Pangenome Reference Consortium (HPRC) [10], the Human Genome Structural Variation Consortium (HGSVC) [11], and the 1000 Chinese Pangenome project [12] have generated high-quality assemblies for population-scale variant discovery and analysis.

However, generating these assemblies typically requires extensive sequencing resources from the target individual (the “donor”), including technologies such as PacBio High Fidelity (HiFi), ultra-long Oxford Nanopore Technologies (UL-ONT), Hi-C, Strand-seq, and Illumina, sometimes supplemented with parental or even grandparental samples [6]. Such extensive data collection is often expensive and inaccessible for individual research groups with limited sequencing budgets.

A more cost-effective and practical alternative is to enhance a draft *de novo* assembly using computational methods such as reference-based scaffolding. In this process, each contig from the draft assembly is aligned to a reference genome to determine its likely position and orientation in the donor genome. The contigs are then ordered and oriented accordingly, typically concatenated with runs of ambiguous bases (Ns), such as the 100 Ns used by RagTag [13]. While reference-based scaffolding reconstructs an approximation of the genome structure, it introduces potential uncertainties. Short contigs, repetitive regions, or common structural variants (such as inversions) can all create ambiguities, meaning that contigs placed next to each other with intervening Ns may not truly be adjacent in the donor genome, and the true sequence spanning the gap remains unknown.

Patching goes beyond scaffolding: it not only determines the order and orientation of the contigs, but also attempts to recover the sequence content that lies between them. In patching, the draft assembly is aligned to a reference genome, and when two contigs are found to be adjacent in the reference, the algorithm fills the gap using the corresponding reference sequence. This approach eliminates ambiguity of filler Ns and produces a more continuous assembly. While the sequence used to patch between contigs may not perfectly match the true donor genome, it is a reasonable approximation, especially when the structure of reference genome is similar to that of the donor, or when the gap being patched belongs to a single haplotype block.

Existing patching tools rely on a single genome to determine contig placement and to supply gap-filling sequence [13, 14]. In this work, we introduce ImpuT2T, a pangenome-based assembly patching tool. Unlike previous methods, ImpuT2T leverages multiple high-quality reference assemblies, not only to infer the optimal order and orientation of contigs, but also to select the most appropriate sequence from available haplotypes to fill the gaps. Our results demonstrate that ImpuT2T outperforms single-reference patching approaches, yielding lower patching error rates, greater recovery of true donor sequence, and higher assembly contiguity. As the number of available T2T quality genomes increases, ImpuT2T offers a cost-effective and accurate way to enhance draft assemblies in the new era of pangenomics. ImpuT2T is implemented as a Snakemake workflow and is available at https://github.com/maojanlin/ImpuT2T.

## 2 Results

### 2.1 Datasets and Materials

In this study, we evaluated the ImpuT2T patching method using four de novo assemblies: HG002, HG005, CN1, and PAN027. These assemblies were generated from donors’ HiFi long reads and parental short reads using the hifiasm assembler [3] with the trio-binning mode (see Section 4.2 for details). In addition, we included another HG002 dataset from the assembly from HPRC Release 1 (HPRC1) [10] to test different quality query assemblies for the same individual. For all experiments, HPRC Release 2 (HPRC2) [15] served as the pangenome reference panel. Since all the individuals in the test set are also in the HPRC2, to avoid bias, we excluded the corresponding haplotypes from the pangenome during each patching experiments. Notably, because T2T-CHM13 utilizes HG002’s Y chromosome as its reference, and T2T-CHM13 was involved in most analyses, we also excluded the Y chromosome of HG002 (both *de novo* and HPRC1) during evaluation.

As benchmarking references, we used high-quality, chromosome-resolved assemblies: Q100 HG002 [5], PAN027 [16], and CN1 [6]. For HG005, since a direct chromosome-resolved assembly was not available, the HPRC2 assembly was scaffolded using YAO [7] with RagTag [13] to create a benchmark.

The quality of patching results was assessed using Genome Quality Checker (GQC) [5], an assembly benchmarking tool. In all analyses, we excluded the short arms of the five human acrocentric chromosomes, as their highly repetitive and complex nature confounds accurate evaluation (see Section 4.4.2 for details).

### 2.2 Overview of the ImpuT2T workflow

ImpuT2T uses a divide-and-conquer approach for pangenome-based patching. First, the input query assembly is compared to a selected set of high-quality (HQ) reference genomes from HPRC2, including T2T-CHM13, GRCh38, HG002, CN1, YAO, and KSA001. The contigs from the query assembly are assigned to chromosomes based on a voting scheme across the HQ reference set (see Section 4.3.2 for details). An optional misassembly correction step can be applied prior to chromosome assignment: large structural variants not present in HQ reference set are removed. After chromosome assignment, contigs belonging to each chromosome are processed independently in a parallelized patching module. The resulting patched chromosomes are then aggregated to produce the final patched genome (Figure 1 **a**).

**Figure 1:**
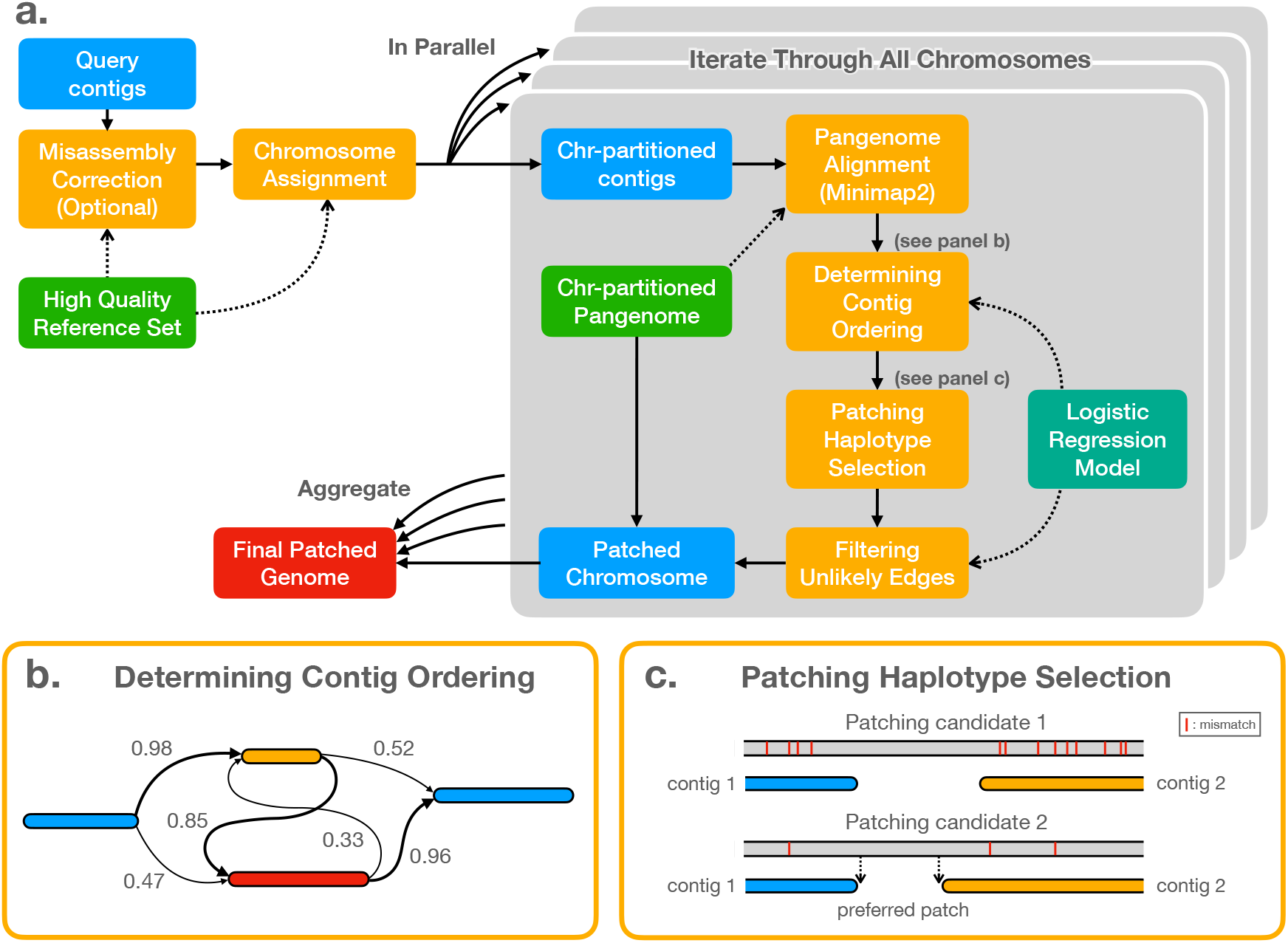
**a**. Pipeline flowchart of ImpuT2T pangenome-based patching method. The “High Quality Reference Set” is composed of T2T-CHM13, GRCh38, HG002, CN1, YAO, and KSA001, total of 10 haplotypes. Solid arrows indicate data transfer, while dashed arrows indicate information flow. **b**. Determination of contig ordering by selecting the graph traversal with the highest aggregated connection probability; edge labels indicate the probability that a given connection belongs to the ground truth. **c**: Selection of a pangenome haplotype to patch a connection. The bottom reference is the preferred choice in this example.

In the patching module, the chromosome-partitioned query contigs are aligned using Minimap2 [17] to a chromosome-partitioned HPRC2 pangenome reference. For each reference haplotype, contigs that align close to each other on the same reference genome are potential neighbors in the scaffolding, and could also be patched together. We consider all such instances as connections (edges) in a patching graph, where nodes correspond to contigs and edges represent candidate connections between them. A pre-trained logistic regression model is then used to estimate the probability that each connection is real. The algorithm searches for a path through the graph that maximizes the total probability of selected connections, as a result determining the optimal order and orientation of the contigs (see Figure 1 **b** and Section 4.3.4 for details).

Once the contig ordering is determined, the algorithm selects a haplotype from the pangenome to patch each gap. This selection is based on the sequence identity of the flanking regions from the two adjacent contigs (Figure 1 **c**). By default, the algorithm evaluates 5, 000 bp from each flank and chooses the haplotype with the highest combined identity score. If there is a tie, the algorithm extends the evaluation to 25, 000 bp from each flank to break the tie.

After determining the patching haplotype for each gap, the algorithm performs a final quality check: any connection with a low prediction score and a patched-to-contig length ratio exceeding a predefined threshold is flagged as low-confidence and filtered out. Following this post-processing step, the remaining patched chromosomes are aggregated to construct the final patched genome. The entire ImpuT2T workflow is organized with the Snakemake workflow.

### 2.3 Quality measurements of a patched genome

We assess the quality of a patched genome using three primary criteria: coverage of the benchmark genome, the number of errors, and contiguity. To evaluate these metrics, the GQC aligns the query genome to the benchmark genome using Minimap2 [17]. The error count includes substitutions, insertions, and deletions (indels). Benchmark coverage is measured as the proportion of the benchmark genome that is mapped by the query genome. For all benchmarks, we exclude the short arms of the five human acrocentric chromosomes from evaluation.

Generally, patching is a trade-off between increasing benchmark coverage and introducing additional errors, as it adds external sequences beyond those present in the donor genome. However, patching does not always result in increased benchmark coverage. For example, in ImpuT2T, some gaps have negative distances, meaning that the adjacent contigs actually overlap. In such cases, the overlapping sequence is trimmed and discarded after patching (Section 4.3.7). Similarly, short contigs that cannot be confidently joined with the main assembly are also excluded (Section 4.3.6).

Contiguity provides a complementary measurement to error count and coverage. While a patched genome may achieve greater coverage, it may still remain fragmented. Although metrics like N50 and NG50 are commonly used to quantify contiguity, we use auN and auNGA, which provide a more contiguous assessment involves the whole Nx curve [18]. The auN is defined as:

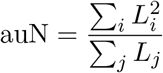

where *L_i_*is the length of the *i*-th contig, and the denominator is the total sum of all contig lengths. auNGA is the auN value normalized by genome length, accounting only for segments that can be aligned contiguously to the benchmark genome, rather than the entirety of each contig.

We report auN for both the benchmark genome and the input query assemblies, as shown in Figure 2 **a** and **b**. For any genome that has been patched, we assess contiguity using auNGA (as in Figure 2 **c**). Although patching can seemingly “merge” two contigs and inflate the contiguity estimate with metrics like auN, incorrect merging will be revealed as breaks in the alignment to the benchmark, meaning such errors will not improve auNGA. In other words, auNGA measures the contiguity contributed only by correctly patched regions.

**Figure 2:**
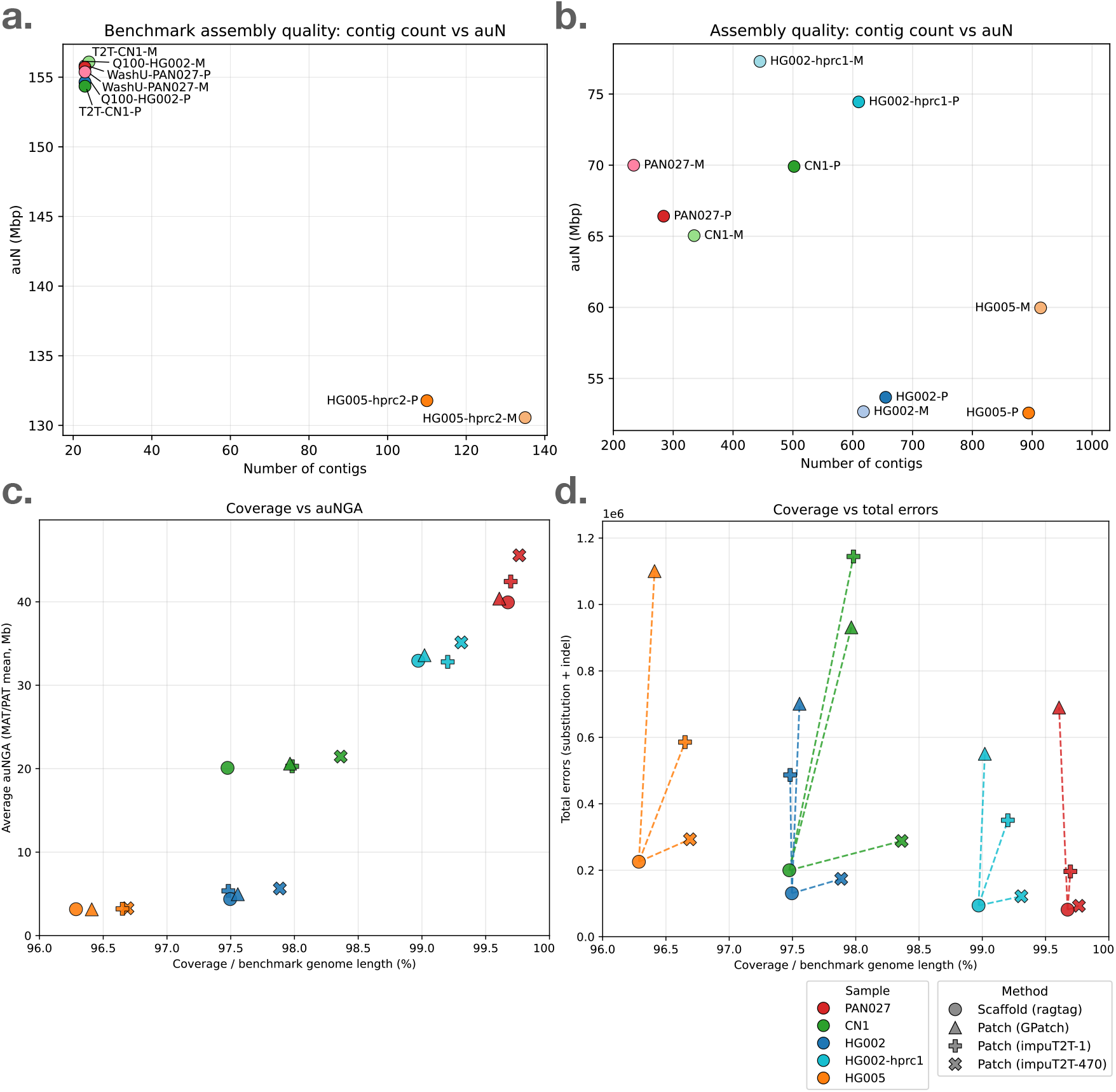
Comparison across the five experimental datasets. **a**: Quality metrics of the benchmark genomes used in this study. P: paternal, M: maternal. **b**: Quality metrics of the four *de novo* assemblies and HPRC1 HG002 genomes used as input queries. P: paternal, M: maternal. **c**: Benchmark genome coverage and auNGA values for each of the five assemblies with different enhancing methods. **d**: Benchmark genome coverage and error counts for each of the five assemblies with different enhancing methods.

#### 2.3.1 Quality of benchmark and input assemblies

Figure 2 **a** presents the benchmark genomes used in our study. HG002, CN1, and PAN027 are haplotype-resolved assemblies with exactly one contiguous sequence representing each chromosome. By contrast, the HG005 assembly from the HPRC2 remains rather fragmented, with each haplotype consisting of more than 100 contigs. Within the HG002, CN1, and PAN027 cluster, the paternal haplotypes of CN1 and HG002 exhibit lower auN values. This is because both paternal haplotypes include chromosome Y, while all the other haplotypes in this cluster carry chromosome X.

Figure 2 **b** depicts the contiguity of the five input query assemblies. HG002 from HPRC1, PAN027, and CN1 display higher contiguity than the *de novo* assemblies of HG002 and HG005. The differences are likely attributable to the variations of HiFi sequencing read quality and sequencing depth (Section 4.2).

#### 2.3.2 Advantages of pangenome over a single reference

To assess the impact of different genome enhancing strategies, we compared the following methods. First, we used RagTag [13] to scaffold the query assemblies as a baseline; however, only contigs successfully scaffolded were retained, unplaced contigs were discarded (Section 4.1). We also employed GPatch [14], a single-reference-based patching tool, using T2T-CHM13 as the reference genome across all five datasets. Additionally, ImpuT2T was run in two modes: using the full pangenome as reference, and using only T2T-CHM13, to highlight the advantage of pangenome over single-reference patching within the ImpuT2T framework.

Figure 2 **c** summarizes the benchmark coverage and assembly contiguity for all five datasets under these different enhancement methods. Pangenome patching led to substantial improvements in genome coverage across all test sets. The gains in contiguity were most pronounced for PAN027, HPRC1 HG002, and CN1, where the initial query assemblies comprised longer contigs. In contrast, the *de novo* HG002 and HG005 assemblies, being more fragmented, exhibited less contiguity improvements following patching.

Although pangenome-based patching methods achieve higher coverage of the benchmark genome, they do not increase the total query sequence length beyond that of single-reference-based patching (Figure S1). Pangenome-patched genomes are more comparct and exhibit fewer unmapped sequence and demonstrate a lower unmapped ratios compared to those generated using single-reference-based patching methods.

Figure 2 **d** presents the benchmark coverage along with total error counts for each method. While error counts in the RagTag scaffold reflect assembly errors, the additional errors in patched genomes represent the expected trade-off of introducing external donor sequences for benchmark coverage. ImpuT2T using CHM13 as the sole reference generally resulted in fewer additional errors than GPatch, except for the CN1 dataset. This may be attributed to ImpuT2T’s more conservative approach to patching connections deemed low-confidence (see Section 4.3.3 for details). The pangenome patching strategy introduced the least amout of error while achieving the highest benchmark coverage.

### 2.4 Performance stratified by chromosome

To build upon our evaluation of overall patched genome quality, we next examined performance at the chromosome level. In figure 3 we directly compare the error counts and benchmark coverage, for each chromosome and haplotype, between genomes patched using ImpuT2T (pangenome-based) and those patched using GPatch (single-reference based).

**Figure 3:**
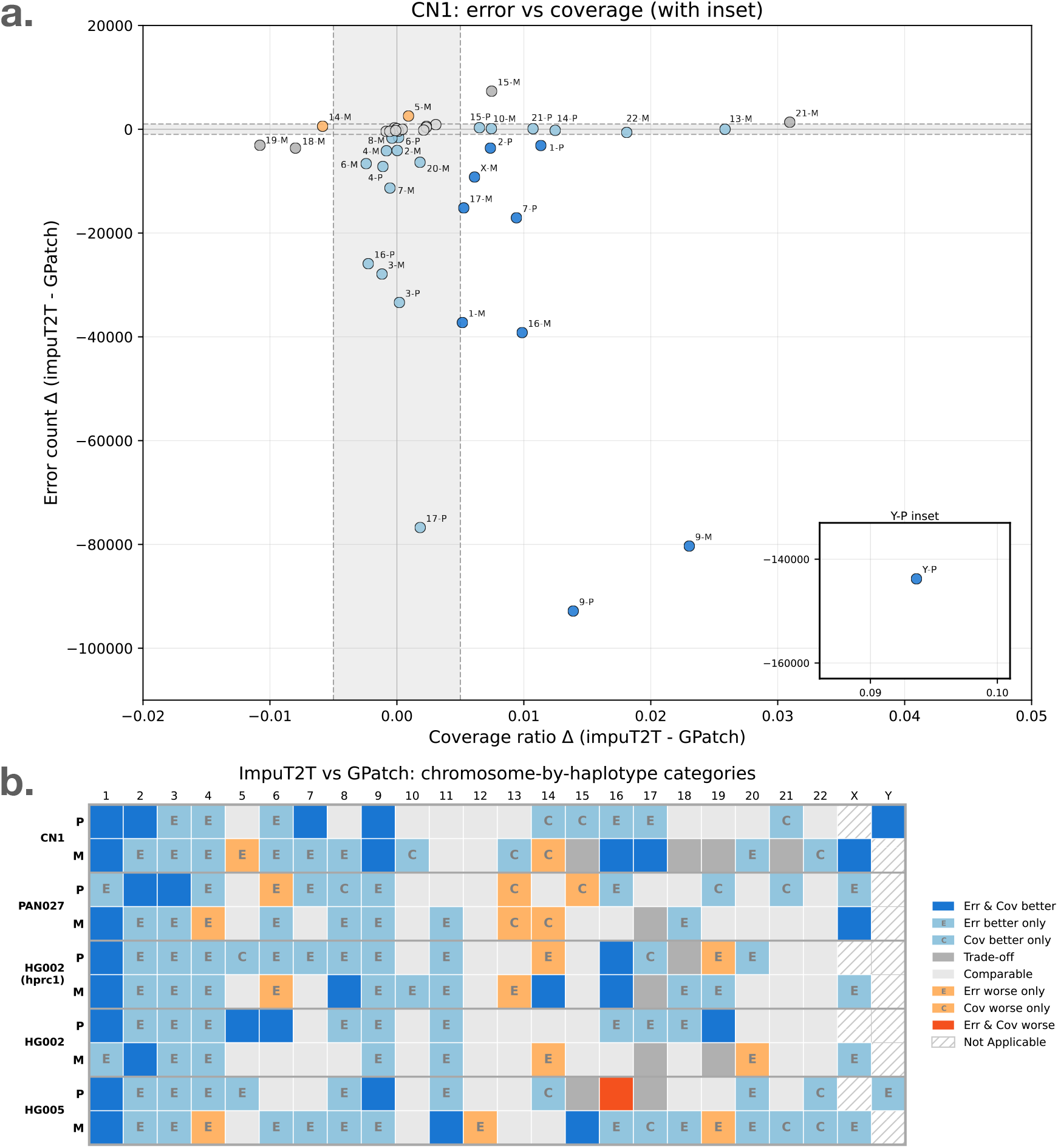
Comparison of error counts and benchmark genome coverage between genomes patched with GPatch and ImpuT2T, stratified by chromosome. **a**: For each chromosome (from both paternal and maternal haplotypes), the difference in error counts and benchmark genome coverage between the two methods is plotted. Each dot represents a chromosome; the crossed grey region delineates cases where the differences in error count are within 1, 000 or coverage differences are within 0.005, indicating comparable performance between the methods. Dots are color-coded to indicate whether ImpuT2T patching yields improvements or deteriorations in one or both metrics compared to GPatch (see legend and panel **b**). Dark blue dots indicate improvement in both error count and coverage, while light blue dots indicate improvement in only one metric. Grey dots represent chromosomes where the two methods yield comparable results or tradeoffs between the two metrics. Dark orange and light orange dots indicate that ImpuT2T performs worse than GPatch in both or only one metric, respectively. To maintain clarity, dots showing comparable performance between GPatch and ImpuT2T are unlabeled due to high density. Other data points are labeled by chromosome and haplotype (where “M” denotes maternal and “P” denotes paternal). Detailed metrics are provided in Table S3, and S2. **b**: For each haplotype and chromosome across the five datasets, the figure displays whether ImpuT2T patching led to improvement or deterioration in benchmark coverage and error count. Chromosomes showing improvement in only one metric are indicated in light blue (for improved) or light orange (for worsened). The letters “E” and “C” indicate which metric has changed: “E” for error count and “C” for benchmark coverage. In both HG002 samples, the Y chromosome was excluded from evaluation.

To facilitate meaningful classification, we considered the two measurements to be comparable when the difference in error count between the two methods was less than 1, 000 and the difference in benchmark coverage was less than 0.5%. Based on this, each chromosome and haplotype combination was assigned to one of eight categories, reflecting whether ImpuT2T showed improvement, deterioration, or comparable results relative to GPatch, across one or both metrics:

- **Err & Cov better**: ImpuT2T improves both error count (fewer errors) and benchmark coverage.
- **Err better only**: ImpuT2T has a lower error count, but similar in coverage.
- **Cov better only**: ImpuT2T has higher benchmark coverage, and similar error count.
- **Trade-off**: ImpuT2T improves one metric but performs worse on the other (e.g., higher coverage but also more errors, or vice versa).
- **Comparable**: Differences between ImpuT2T and GPatch fall within the predefined thresholds for both error count and coverage, indicating similar performance.
- **Err worse only**: ImpuT2T results in a higher error count, but coverage is similar.
- **Cov worse only**: ImpuT2T results in lower coverage, but error count is similar.
- **Err & Cov worse**: ImpuT2T performs worse than GPatch in both error count and benchmark coverage.

Figure 3 **a** presents a detailed comparison between ImpuT2T and GPatch for all 46 chromosomes of individual CN1, including the 44 autosomes as well as chromosomes X and Y. Among these, ImpuT2T improved both error count and benchmark coverage in 10 chromosomes. In 18 chromosomes, ImpuT2T showed improvement in either error count or coverage, but not both. For 16 chromosomes, the results were either comparable between the two methods or indicated a tradeoff between error count and coverage. Notably, only two chromosomes showed worse results for ImpuT2T in one of the metrics, and none exhibited deterioration in both error count and coverage.

Another important metric is the number of fully contiguous, or so called “T2T”, chromosomes obtained after patching. By design, GPatch generates a single patched contig for each chromosome per haplotype. In contrast, ImpuT2T yields 27*/*46, 33*/*46, 31*/*45, 30*/*45, and 21*/*46 fully contiguous patched chromosomes for the datasets CN1, PAN027, HG002-HPRC1, HG002, and HG005, respectively (Figure S2 **a**). Notably, the number of these single-contig chromosomes produced by ImpuT2T remains relatively stable regardless of pangenome size (Figure S2 **b**). This is because ImpuT2T only fills gaps when there is sufficient confidence (Section 4.3.3). ImpuT2T assumes that the initial set of assembled contigs, after they have been split in the misassemble correction step, are then fully correct, and does not break contigs during the main patching process. Consequently, a misassembled contig, such as one containing an inversion or a chimeric join, can prevent remaining contigs from establishing a true chromosomal connection.

Figure 3 **b** presents a categorical summary across all chromosomes from the 10 haplotypes across our five test datasets. Of the 228 chromosomes evaluated, 112 (53.5%) demonstrated clear improvement in at least one metric (either reduced errors, increased benchmark coverage, or both) when using ImpuT2T. Another 88 chromosomes (38.6%) exhibited either comparable results or a trade-off between error and coverage. Notably, in only 18 cases (7.9%) did ImpuT2T result in deterioration in one or both metrics compared to GPatch. Further details can be found in Table S1.

In most of the improvement cases (76 out of 112), the gain observed with ImpuT2T over GPatch is primarily in the reduction of error count, rather than increasing of benchmark coverage. This typically occurs when both the single-reference (T2T-CHM13) and the haplotype selected by ImpuT2T from pangenome arrange contigs in the same order, but the selected haplotype aligns more accurately with the benchmark, resulting in fewer errors. However, because the overall structure matches, the increase in benchmark coverage is minimal.

In contrast, improvements in benchmark coverage usually indicate that ImpuT2T’s chosen contig ordering more closely matches the benchmark sequence, or that there are large-scale discrepancies between the single-reference and the benchmark. For example, if the benchmark contains a long insertion but the two contigs overlap on the single-reference, the entire inserted sequence can be missed by the single-reference-based patching algorithm, resulting in substantially lower coverage.

Notably, ImpuT2T consistently achieves substantial improvements on chromosomes 1 and 9 across all datasets and haplotypes. This likely reflects the unique genomic features of these chromosomes: both chromosome 9 and 1 have the largest pericentromeric regions in the human genome [19]. In addition, chromosome 9 contains the longest human satellite array and is highly polymorphic [20]. These characteristics make patching these long, repetitive pericentromeric regions with only a single reference genome particularly error-prone, providing an opportunity for a pangenome-based patching strategy to improve the result.

On the other hand, for chromosome 12, both single-reference-based and pangenome-based patching methods generally yield comparable results across samples. Detailed examination of the error counts and coverage (see Table S2 and S3) reveals that both Im-puT2T and GPatch perform well on this chromosome, with minimal increases in error count relative to the scaffold (raw contig) assemblies. This is likely because the input contigs for chromosome 12 are already of high quality, leaving little room for further improvement.

### 2.5 Impact of pangenome composition

In the previous sections, all pangenome patching experiments utilized the full HPRC2 panel after excluding the donor’s paternal and maternal haplotypes. To better understand how pangenome composition influences patching performance, we evaluated the impact of pangenome size and population specificity. Haplotypes in the HPRC2 panel were classified into five superpopulations, AFR, EAS, EUR, AMR, and SAS, according to the 1000 Genomes Project nomenclature from the International Genome Sample Resource (IGSR) (see Section 4.4.1 for details).

We constructed two types of subset pangenomes from HPRC2: balanced and population-specific. In all cases, T2T-CHM13 and GRCh38 haplotypes were always included as the first two entries in each subset pangenome (with T2T-CHM13 designated as EUR and GRCh38 as AFR). For the balanced subsets, haplotypes were selected in a round-robin fashion from the five superpopulations to ensure even representation, continuing until one group was exhausted, and then cycling through the remaining superpopulations. We generated balanced subsets containing 1, 5, 10, 25, 50, 100, 200, 300, and up to all 470 haplotypes. Because the EUR superpopulation contains the fewest haplotypes (65, including T2T-CHM13), 300 is the largest subset size for which a balanced distribution across all superpopulations can be maintained. In contrast, population-specific subsets included only haplotypes from a single superpopulation matching the ancestry of the test individual. We focused on AFR, EUR, and EAS populations, as these correspond to the ancestries in our benchmarking datasets.

For the EAS and AFR superpopulations, we tested subsets containing 10, 25, 50, and 100 haplotypes, reflecting the approximately 100 available haplotypes per group. For EUR-specific subsets, due to the smaller number of available haplotypes, we evaluated subsets of 10, 25, 50, and 64 haplotypes (the maximum available). It is important to note that both T2T-CHM13 and GRCh38 were also included in the population-specific panels, even though they may not belong to the same superpopulation as the test sample.

Figure 4 **a** and **b** illustrate the impact of subset pangenome composition on CN1, presenting error count, benchmark coverage, and auNGA across different subsets. For CN1, EAS represents the matching population and AFR is used as an outgroup. In these plots, the point labeled “1” corresponds to patching with only T2T-CHM13, while “P” indicates patching using the benchmark CN1 genome. Both the balanced and EAS-specific subsets show a trend where increasing the number of haplotypes leads to greater coverage, fewer errors, and higher auNGA values in the patched genome. However, this trend is not strictly monotonic. For instance, within the balanced subset, the 50-haplotype panel yields higher coverage than the 100- and 200-haplotype subsets. Likewise, in the EAS-specific subset, the 50-haplotype panel produces more contiguous assemblies (higher auNGA) than the 100-haplotype panel.

**Figure 4:**
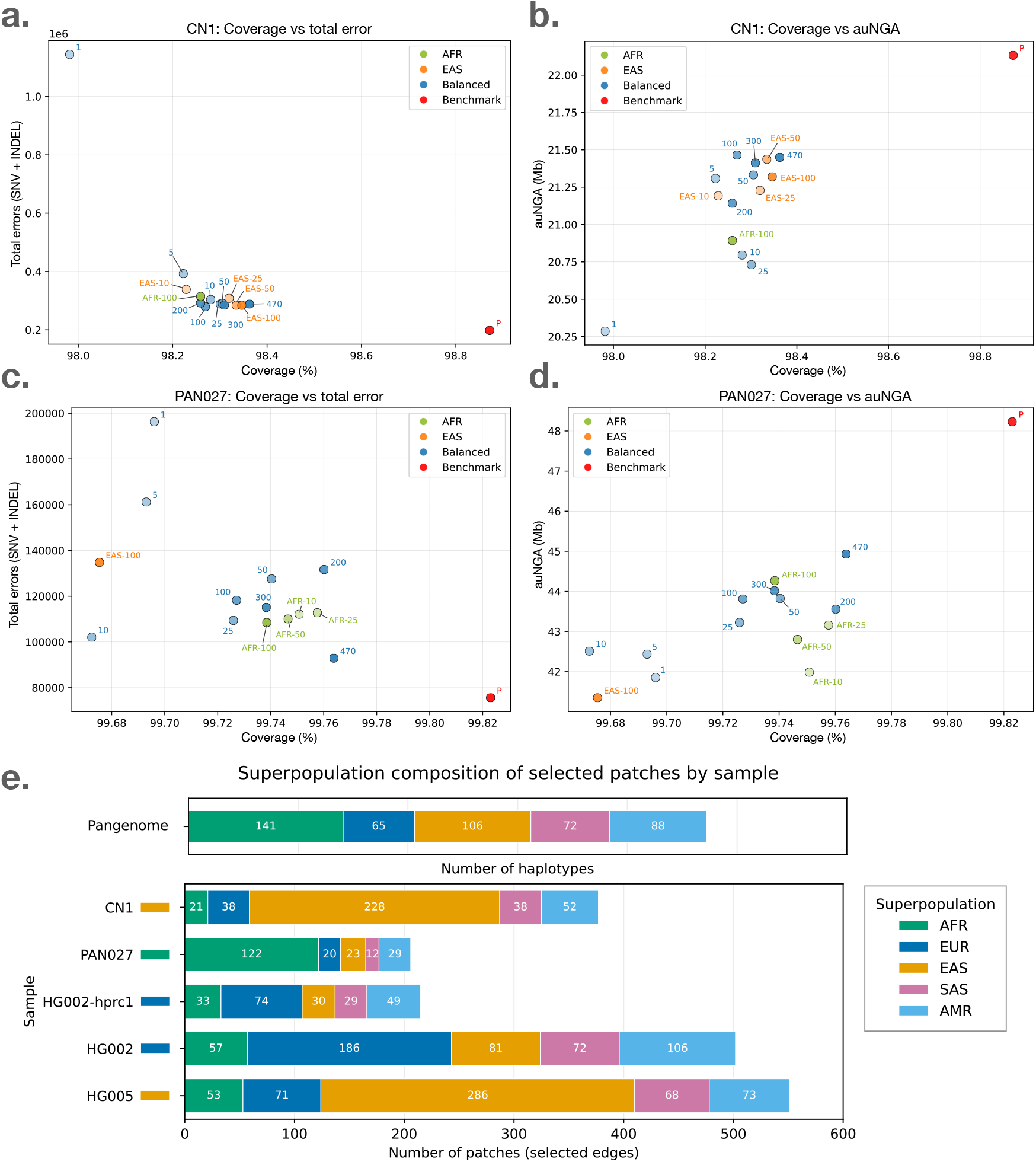
Impact of pangenome size and population-specific pangenomes on ImpuT2T performance. **a** and **b**: Benchmark genome coverage, total error count, and auNGA for CN1 using pangenomes of varying sizes (5 to 470 haplotypes). EAS denotes the population-specific pangenome matching CN1. **c** and **d**: Benchmark genome coverage, total error count, and auNGA for PAN027 across the same pangenome size range. AFR denotes the population-specific pangenome matching PAN027. **e**: Stacked bar plot illustrating the superpopulation origins of patched sequences for each sample when using the full pangenome (470 haplotypes). The top panel displays the superpopulation composition for the HPRC2 pangenome. For the patched dataset in the bottom panel, colored bars indicate the proportion of patches derived from each superpopulation, with a color label indicating the matched superpopulation for each sample.

Figure 4 **c** and **d** show the performance of different subset pangenomes on PAN027, where AFR is the matching population and EAS is the outgroup. Here, the differences in benchmark coverage between the different subsets are smaller compared to CN1. Nevertheless, the overall trend persists: subsets with more haplotypes generally achieve better results. In both CN1 and PAN027, using a population-specific pangenome from an outgroup leads to poor performance. While the matching population-specific pangenome does not always outperform the full panel of 470 haplotypes, it appears to be more efficient; for example, a 100-haplotype subset from the matching population performs comparably to a 300-haplotype balanced subset. Similar trends are seen in the HG002 dataset using both *de novo* assembly and HPRC1 input, as well as in HG005 (see Figure S3–S6).

Figure 4 **e** shows the superpopulation composition of all HPRC2 haplotypes in the top row. The following five rows present the superpopulation origin of each ImpuT2T patch (gap) for the respective dataset when using the full HPRC2 panel. Each number in these rows corresponds to a patched gap, including instances where the gap is negative (i.e., adjacent contigs overlap). For every dataset, the plurality of the patches originate from haplotypes belonging to the superpopulation that matches the sample’s ancestry: EAS for CN1 and HG005; AFR for PAN027; and EUR for both HG002 assemblies. There is also a clear inverse relationship between the number of patches and the input assembly quality: the highest-quality assemblies, PAN027 and HG002-HPRC1, require the fewest patches; CN1 is intermediate; and the *de novo* assemblies of HG002 and HG005 require the most.

### 2.6 Time and memory usage

Figure 5 summarizes the runtime and memory consumption of ImpuT2T using pangenomes of different sizes. For comparison, we include measurements for CN1 *de novo* assembly and GPatch patching of CN1 using T2T-CHM13. The *de novo* assembly times reflect only the core trio-binning step, excluding the yak preparation procedure. All performance tests were conducted using compute nodes with 48 threads. Panels **a** and **b** display the elapsed wall-clock time and total CPU hours across the different methods. In panel **b**, the ImpuT2T pipeline is further broken into four stages: Prepare, Align, Local, and Patch & final. The Prepare stage corresponds to chromosome assignment as described in Figure 1. The Align stage aligns chromosome-partitioned contigs to the relevant pangenome using Minimap2 [17]. In the Local stage, flanking sequences around each contig are aligned to the pangenome to determine flanking score. The Patch & final stage includes contig ordering, haplotype patch selection, filtering, and construction of the final patched genome.

**Figure 5:**
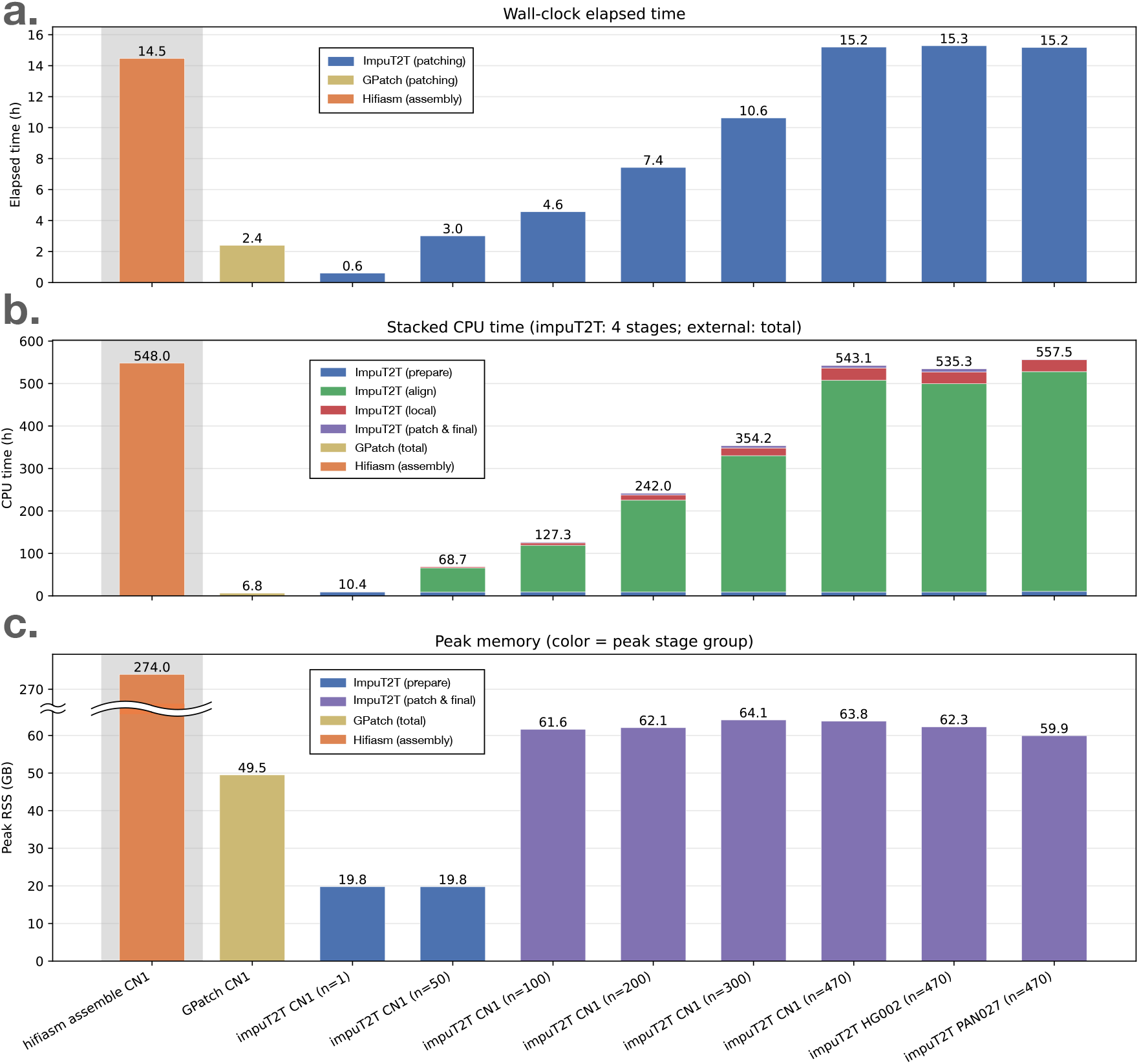
Time and memory usage of the patching methods. Results for *de novo* assembly using hifiasm trio-binning are also shown for comparison; only the assembly step is measured, excluding k-mer counting with yak. All experiments were run using 48 threads. **a** and **b**: Elapsed wall-clock time and total CPU hours for GPatch and ImpuT2T with different pangenome sizes. We measure CN1 with 1, 50, 100, 200, 300, and all available haplotypes. For HG002 and PAN027, only the full pangenome was measured. **c**: Peak memory usage for each patching method. For ImpuT2T, color indicates the pipeline stage with the highest memory consumption.

The majority of ImpuT2T’s total runtime is spent in the Align stage, which is highly parallelizable. Consequently, the ratio of CPU hours to wall-clock time is roughly 35 when utilizing 48 compute threads. Notably, ImpuT2T’s elapsed time for a 50-haplotype pangenome is comparable to that of GPatch using a single reference. While GPatch also relies on Minimap2 for alignment, it performs an all-against-all alignment of contigs to T2T-CHM13, whereas ImpuT2T first assigns contigs to chromosomes and aligns them within these partitions. As a result, for single-reference patching, ImpuT2T has greater CPU time than GPatch—primarily due to its preparatory chromosome assignment stage—whereas for larger pangenomes most of the runtime shifts to the highly scalable Align stage.

In addition to CN1, we evaluated the runtime and memory usage for HG002 and PAN027 using the full HPRC2 pangenome. All three datasets showed highly similar performance, both in terms of elapsed time, CPU usage, and memory footprint. Regarding memory (panel **c**), ImpuT2T with the full panel uses slightly more memory than GPatch, but the requirements remain comparable. Both patching methods require only a fraction of the memory needed for hifiasm trio-binning *de novo* assembly. Thus, any computing environment capable of running hifiasm assembly can comfortably execute ImpuT2T with the full HPRC2 pangenome.

## 3 Discussion

Recent advances in long read sequencing technology and the releases of high-quality, near T2T assemblies have made it feasible to use pangenome resources for lower-quality assemblies enhancement. ImpuT2T is designed to bridge the gap between the resource intensive state-of-the-art assembly methods and what can be easily achieved in a typical laboratory setting.

Our results demonstrate that ImpuT2T, utilizing a pangenome reference panel consistently produces assemblies with lower error rates, greater recovery of donor sequences, and higher contiguity compared to patching using a single reference genome. We evaluated the effect of different pangenome sizes as well as population-specific reference panels. Both balanced and population-specific panels exhibited a clear trend: as the number of haplotypes in the reference increased, the patched genome better resembled the benchmark sequence. Notably, performance did not plateau even at the full panel size of 470 haplotypes.

We also observed that using a population-specific pangenome matching the donor’s superpopulation could achieve a similar level of performance with fewer haplotypes than a balanced, multi-population panel. However, in our experiments, the full pangenome panel usually outperformed or matched the best results from population-specific subsets. Therefore, we recommend utilizing the complete panel whenever computational resources permit, as this both maximizes recovery and mitigates risks related to incorrect or ambiguous donor ancestry—for example, in cases of admixture or uncertain population assignment.

We benchmarked ImpuT2T against the single-reference-based method GPatch [14], demonstrating the benefits of utilizing pangenome information. We also considered RagTag patch mode [13], but consistent with previous reports [14], we found that RagTag Patch mode is unreliable for human genomes. RagTag failed in many of our trials. Accordingly, we did not include results for RagTag patch in our main analyses, using only RagTag scaffold mode as a point of reference. Another notable tool is panpatch [16], but this method is designed for situations where multiple assemblies of the same individual or pedigree-based reference pools are available, and requires a user-specified patching order. Because its application and goals differ from those addressed by ImpuT2T, we did not include panpatch in our primary evaluation (see Section 4.1 for further discussion).

One crucial limitation of ImpuT2T is its time complexity and scalability. While the workflow is highly parallelizable and can be completed in a runtime and with peak memory usage comparable to *de novo* assembly using hifiasm, scaling to increasingly larger pangenome panels presents challenges as more phased assemblies become available. In practical terms, any compute node capable of assembling a *de novo* assembly should also be able to perform patching with the full HPRC2 reference set using ImpuT2T. However, as the size of the reference panel continues to rise, further improvements in scalability will be necessary. One potential solution is to adopt r-index-based alignment methods, which allow indexing the entire pangenome and performing alignment in one single query [21].

Alternatively, graph genome aligners such as Giraffe for long reads [22] may offer a scalable solution. However, these approaches have their own tradeoffs: the underlying graph indices can be extremely large, and workflows are not always straightforward to deploy. To address this, some researchers have constructed “personalized” pangenome graphs by subsampling to a limited number of haplotypes [22, 23]. The complex repetitive regions are often masked out when building a pangenome graph index [24]. In contrast, the ImpuT2T framework’s strategy of iterating through linear genome collections offers greater flexibility and full resolution in complex genomic regions.

There are several areas where the ImpuT2T workflow, which balance performance and efficiency, could be improved. One key design decision is that the workflow involves initially assigning contigs to chromosomes based on their similarity to a high-quality (HQ) reference assembly, after which each chromosome undergoes patching independently. This strategy helps modularize the patching process and reduces computational complexity compared to an all-to-all comparison. However, errors in the initial contig assignment can propagate through the later stages. A misplaced contig not only leaves a gap in the donor chromosome, but also leads to suboptimal results in the assigned chromosome. Particularly, we observed that the assignment of contigs originating from acrocentric short arms is usually unreliable. Because these regions are highly repetitive, especially the ribosomal DNA arrays, and because GQC relies on Minimap2 alignments that are less consistent in such sequences, we excluded them from all GQC-based evaluations (Section 4.4.2). Furthermore, if contigs are misassembled (e.g., a chimeric contig that mistakenly spans two different chromosomes) or the donor carries true structural rearrangements such as Robertsonian translocations, the current framework can be ineffective due to the reference pool being limited to one specific chromosome.

Another aspect of the workflow is the optional misassembly correction step that can be applied prior to running the main workflow. The correction step remove the connection discordant to all HQ haplotypesthe. However, this correction is a double-edged sword: while it can successfully break apart erroneous misjoints, it also can potentially remove private structural variants not in the HQ set. Therefore, we recommend using this correction step selectively, mainly when the input assembly quality is known to be low.

Both the contig-to-chromosome assignment step and the optional misassembly correction rely on the HQ reference set, which is relatively small and not fully representative of global human population diversity. Looking ahead, as more T2T-level assemblies become available, it will be possible to construct a more robust and population-representative HQ reference set, which should improve these aspects of the ImpuT2T workflow.

## 4 Methods

### 4.1 Running existing tools

#### GPatch

We evaluated GPatch version 0.4.0 in our experiments, using T2T-CHM13 [4] as the patching reference across all tests. For consistency, the input contigs provided to GPatch matched those used in the primary ImpuT2T workflow. For HG002 and HG005, we used the “corrected” assemblies in both ImpuT2T and GPatch. Since hifiasm produces phased assemblies, we ran GPatch separately for each haplotype, paternal and maternal, per sample. In cases where a given haplotype assembly was not expected to contain one of the sex chromosomes, any GPatch output corresponding to the absent chromosome was excluded from further analysis.

#### Panpatch

Panpatch is primarily designed to recursively patch a donor genome against different assemblies of the same individual (for example, generated using different sequencing platforms) or against closely related assemblies from the same pedigree. While panpatch utilizes a pangenome graph to facilitate patching, it selects patch sequences based on a user-defined order rather than automatically identifying the best-fitting sequence through sequence content comparison. This approach is well-suited to situations where multiple assemblies of the donor or parental genomes are available with varying quality. However, it differs from approaches typically used in broader pangenome contexts. In our testing on a small pangenome comprising 25 haplotypes, we observed that most patches were derived from just the first few references specified. The first individual in the list accounted for approximately 75 80% of all patches in all of our experiments. Based on these observations, we concluded that panpatch operates under a distinct paradigm compared to methods designed for large-scale pangenome patching.

#### RagTag

In all of our experiments and throughout the ImpuT2T workflow, we used RagTag v2.1.0. For benchmarking different assembly improvement strategies, we based our comparisons on the output from `ragtag.py scaffold`. RagTag `scaffold` produces an `.agp` file detailing the scaffold structure and which contigs are assigned to which chromosomes of the reference (T2T-CHM13 in our experiments). During evaluation, we retained only the scaffolded contigs and excluded any unplaced contigs, as these unplaced sequences are generally difficult to use in practical settings.

We also performed a brief assessment of `ragtag.py patch`. We observed that specifying the `--aligner minimap2` option increased the likelihood of RagTag `patch` producing outputs compared to the default `--aligner nucmer` setting. However, Rag-Tag `patch` still fail to generate patches in many cases, consistent with previous reports [14]. For instance, in our CN1 tests, RagTag `patch` successfully produced a patch for chromosome 20 using T2T-CHM13, but failed to patch chromosome 1 in the same setup. Therefore, we did not include RagTag `patch` in our main comparisons and used only the RagTag `scaffold` results as a reference.

### 4.2 *de novo* assembly

All four *de novo* assemblies were generated using hifiasm (trio-binning mode, version 0.18.2-r467) [3]. HiFi reads for CN1 and PAN027 were obtained directly from their source repositories. For HG002 and HG005, HiFi reads were extracted from their GRCh38-aligned BAM files using `samtools fastq`. Parental short-read data were processed with `yak count -k31 -b37 -t46` using recommended parameters. All hifiasm runs utilized 48 threads.

Sequencing statistics for the donor HiFi and parental short-read data used in these assemblies are summarized in Table S4. Parental short reads for CN1 were generated on the DNBSEQ platform, whereas those for HG002, HG005, and PAN027 were generated on Illumina. HiFi coverage ranged from 35.7*×* (HG002) to 58.8*×* (CN1). Among the four datasets, CN1 had the highest HiFi coverage, while PAN027 had the longest HiFi read N50.

### 4.3 ImpuT2T workflow

#### 4.3.1 Assembly correction: Mumento correct

`Mumemto correct` operates as an independent program designed to improve assembly accuracy. It compares input contigs against the high-quality (HQ) haplotype set by identifying all maximum unique matches (MUMs) using Mumemto [25], which are then stored in a bumbl file. To optimize performance, the MUM record for the HQ set is precomputed; the target contigs can be efficiently incorporated using `Mumemto merge`. Once the MUM record is established, the method scans the MUMs along each contig. If two colinear MUMs on a contig exhibit a discordance, i.e. the distance between them differs from that in all HQ haplotypes by at least 200 kb, this indicates a potential misassembly. The contig is split at the discordant place only if the discordance persists across five consecutive MUMs, thereby reducing the chance of splitting due to random or isolated errors. Each split breakpoint is then trimmed by 3 kb on either side to ensure clean boundaries.

Highly variable regions of the human genome, such as centromeres and acrocentric short arms, can differ greatly between individuals. As a result, these regions are excluded from the correction process. Specifically, if either of the discordant colinear MUMs falls within a highly variable region in CHM13, the discordance is ignored and no contig split is performed at that site.

While this correction step effectively removes misassemblies, it may also eliminate private structural variants absent from the HQ set. Therefore, we recommend applying it primarily to low-quality assemblies.

#### 4.3.2 Chromosome assignment

The ImpuT2T workflow begins by running RagTag scaffold [13] on the input contigs (or split contigs, if the correction step was applied) using all HQ haplotypes as references. For each haplotype scaffold assignment, a contig-to-chromosome assignment is considered valid if the contig is sufficiently long (*>* 100 kb), or if at least two different haplotypes assign the same contig to the same chromosome. These criteria help filter out spurious assignments, particularly those arising from short contigs with repetitive sequence.

After this initial assignment, it is possible for a given contig to be assigned to multiple chromosomes. To address this, we assess whether there is strong consensus among the HQ haplotypes for a particular chromosome assignment. For example, if the majority of HQ haplotypes support placing a contig on a specific chromosome and there are only one or very few disagreeing assignments, we retain only the majority assignment. Practically, for each contig, we identify the chromosome with the highest number of supporting assignments, and retain any chromosome assignment with support greater than half of this maximum. Only in this situation can a contig be used in patching multiple chromosomes.

#### 4.3.3 Building connections from Pangenome alignment

After chromosome assignment, ImpuT2T call `minimap2 -x asm5` to align all the contigs to their corresponding chromosome-partitioned pangenome. The alignment result is in `paf` format and passed down to further analysis if the contigs show adjacency according to the corresponding reference.

In the alignment process, a single contig may be fragmented into multiple aligned segments, and in some cases, a given sequence can map to several locations within the reference. To ensure quality, ImpuT2T considers an aligned segment valid only if it meets the length criteria: the segment must constitute at least 5% of the total contig length or be longer than 10, 000 base pairs. In all following analyses, only these valid segments were considered.

A segment end is considered “connectable” only if it overlaps the terminal 10% of either end of the contig. When multiple segment ends fall within this terminal region on the same contig end, only the outermost segment (i.e., the segment extending furthest toward the contig boundary) is retained as connectable. For example, in Figure S7, segment ends **a** and **e** are considered “connectable”, whereas **b**, **c**, **d** and **f** are not. This approach operates under the assumption that input contigs are correctly assembled up to their boundaries. In contrast, GPatch [14] allows connections at every segment ends, resulting in a “contiguous” single patch for every chromosome. However, restricting connections to contig ends yields joins that more accurately reflect the structure in the source assemblies.

To reduce redundancy, we discard any alignment contig that is fully contained within another segment, as it is unlikely to provide additional information. A more complex scenario arises when there is an “in-between” alignment located between two split alignments of the same contig. If the two split alignments are oriented in the same direction, we treat them as a single continuous alignment and remove the in-between contig, as illustrated in Figure S8, case 1. Conversely, if the split alignments are in opposite orientations, we retain the in-between alignment, since it may capture meaningful structural variation (see Figure S8, case 2). Note that the in-between contig may itself consist of several split alignments; however, it is considered fully contained only if all these split segments are aligned within the outer alignments.

A connection between two contig segments can be established when their respective connectable ends are oriented toward each other. Specifically, with respect to their alignment positions on the reference, the segment aligned upstream must be connectable at its right end, while the segment aligned downstream must be connectable at its left end, with each segment belonging to a different contig. Candidate connectable segment ends can be classified into three categories: disjoint, overlapping, or contained overlap (see Figure S9). In the case of a contained overlap, the “contained” segment must be part of a contig that extends beyond the boundaries of the overlapping segment. Note that this “contained” segment must originate from a larger contig in which the remaining sequence is not already contained within the same outer contig; otherwise, such a contig would have been discarded during the earlier filtering step.

Disjoint and overlapping connections are considered high-quality, while “contained overlap” connections are acceptable but not preferred. When multiple candidate connections exist, we prioritize them by sorting based on the absolute distance between the two ends on the reference. If the candidate with the smallest absolute distance is disjoint or overlap, we retain only these and discard all “contained overlap” candidates as well as any candidates with an absolute distance greater than 100 kbp. However, if the closest candidate is a “contained overlap”, we retain all available candidates. Note that “contained overlap” connections are also subject to an intrinsic overlap distance constraint relative to the length of the contained contig described in previous section.

##### flanking identity score

Every connection has a flanking score. ImpuT2T performs a terminal re-alignment for the two ends of a contig, and aggregate the two terminal’s alignment score as the flanking score of a connection. Since not all contig are aligned to the end on the reference, the terminal selection for re-alignment is adjusted accordingly.

Figure S10 illustrates how flanking sequences are selected for terminal alignment. Without loss of generality, we define the selection range on the left flanking region. The anchor point is defined as the end of the aligned segment closest to the contig boundary. By default, the terminal length (TL) is set to 5, 000 bp, but is increased to 25, 000 bp in tie-break scenarios (see Section 4.3.5). Around the anchor, a symmetric window of 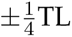 is defined within the contig. For the left end, the selected window extends from 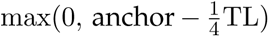 to 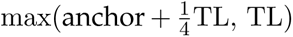. The right flanking region is defined in the same way, but applied at the right end of the contig.

A terminal re-alignment is considered a hit if it falls within 10 kb of the original alignment position. If multiple hits are identified for the same contig end, the one closest to the original position is chosen. If no hit is detected, a score of zero is assigned for that contig end.

#### 4.3.4 Determining contig ordering and orientation

The connection data is aggregated across the pangenome to construct a contig-end graph. Each contig end is represented as a node. Intra-contig edges connect the two ends of the same contig (begin to end), providing fixed links that allow traversal through an individual contig. Inter-contig edges connect ends of different contigs and represent candidate connections between contigs. When multiple pangenome haplotypes support the same connection between two contig ends, these connections are merged into a single inter-contig edge. Each inter-contig connection is assigned a weight equal to the estimated probability that it is true, as predicted by a logistic regression model trained on benchmark data. After an optimal layout is identified by graph search, the resulting traversal specifies both the order and orientation of contigs in the patched chromosome assembly.

##### Logistic regression model

The logistic regression model is trained using the HG002 dataset. Specifically, the benchmark HG002 assembly serves as the reference to patch the *de novo* assembled HG002 contigs, establishing ground truth order and orientation for model training. The model is composed of two features: the number of haplotypes supporting the connection and the highest flanking sequence identity score among all supporting haplotypes. Figure S11 **a** illustrates the distribution of prediction scores for true and false connections in the HG002 training set, while panel **b** shows corresponding results for the CN1 test set. The Receiver Operating Characteristic (ROC) curves for both training and test sets are shown in panels **c** and **d**, respectively. Figure S12 shows additional confusion matrices and test results for PAN027, HG005, and HG002 HPRC1 datasets. The model demonstrates consistently high performance across all test datasets, achieving area under the curve (AUC) values above 0.9. We also evaluated the model on CN1 using pangenomes of different sizes, from the full set of 470 haplotypes down to just 25 haplotypes (Figure S13), finding that predictive performance remains robust even with much smaller pangenomes.

##### Connection graph layout

The goal of the graph search algorithm is to identify a layout consisting of one or more contig chains that visits each contig-end node at most once. Ideally, the layout is contiguous and composed of high-weight inter-contig connections. The algorithm is composed of two stages. In the first stage, a restricted subgraph is constructed from high-confidence inter-contig edges, and a depth-first search (DFS) seeks the layout that maximizes the combined layout score. The DFS stage is subject to a timeout (default: 300 seconds). In the second stage, a greedy procedure follows: it is initialized with the best DFS layout found in the first stage, then processes the remaining inter-contig connections in descending order of edge weight, adding a connection only if both endpoints are unused and the edge weight is at least 0.05.

In the first stage, low-confidence inter-contig edges are excluded before DFS. Any edge with weight below 0.6 is removed. For each contig-end node, edges connected to that node with weight less than one-half of the maximum edge weight at that node are also removed. The remaining candidates are ranked globally by confidence, and only the 20 highest-ranking inter-contig edges are retained for graph search.

Because it is not always possible to connect the entire graph into a single path, we employ a combined scoring scheme to rank the resulting path components (i.e., connected subgraphs or path fragments). For each contig *c*, define the **normalized contig length**

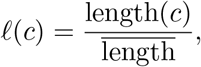

where 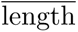 is the mean contig length in the assigned chromosome assembly. For each chain (path fragment) *i*, let *w_j_* denote the aggregated edge weights of inter-contig connection *j*. The chain score is

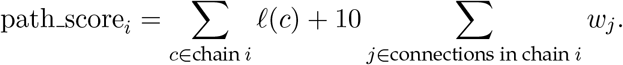

And the overall layout score is

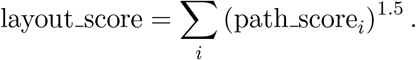

The chain score balances inter-contig connection weights and contig length. As a result, a layout consisting of many short contigs with numerous inter-contig connections is not automatically preferred over one dominated by a long contig with few or no connections. The overall layout score sums (path score)^1.5^ across chains. Raising each chain score to the power of 1.5 favors layouts with fewer, stronger chains over the same total evidence split across many separate fragments.

In the second stage, a greedy procedure extends the best DFS layout from the first stage. Several stage 1 filters are relaxed: the global weight cutoff of 0.6 and the per-node half-max filter are not applied, and up to 200 inter-contig connections (ranked by edge weight) are reconsidered as candidates. Because long-distance connections are less reliable, any connection with distance greater than 2 Mb and edge weight less than 0.5 are excluded; shorter connections are not subject to this requirement. Candidate connections are processed in descending order of edge weight: a connection is added only if both endpoints are unused and the edge weight is at least 0.05. The final contig order and orientation are taken from this extended layout.

#### 4.3.5 Haplotype selection

For each inter-contig connection selected in the layout, multiple pangenome haplotypes may provide support. To determine the most appropriate donor haplotype, we first evaluate the flanking sequence identity scores, based on 5, 000 bp of sequence at each contig end. The score for each end is calculated as the number of residue matches reported by `Minimap2` (i.e., the count of matched bases), so the maximum possible combined score for a connection is 10, 000. We classify the connection as high-confidence if the highest flanking identity score for that connection exceeds 6, 000; otherwise, the connection is considered low-confidence.

For low-confidence connections (maximum score ≤ 6, 000), we restrict consideration to the ten highest-scoring haplotypes, as well as any others with scores at least 50% of the maximum. Among this pool, we select the haplotype with the smallest absolute patching distance.

For high-confidence cases (maximum score *>* 6, 000), we determine whether a single haplotype is clearly better. If the top scorer exceeds the second-best by more than 3 points, that donor is selected. If the score difference is 3 or less, or if multiple haplotypes tie for top score, a tie-break is required. When there is a unique top scorer, the tie set comprises that haplotype together with all haplotypes tied for second place. When multiple haplotypes share the top score, only those top scorers proceed to the tiebreak. If all tied haplotypes report a negative distance (i.e., overlapping contigs), we resolve the tie without additional alignment: the unique top scorer is taken when present, otherwise we choose the top-scoring haplotype whose distance is closest to the mean distance among top scorers.

To break ties, we perform a second round of terminal alignment, this time extending to 25, 000 bp at each contig end and select the haplotype with the highest refined score. It is important to note that not all connections are retained in the final layout, so the tie-breaking procedure is applied only after the final layout has been determined.

Figure S14 **a** illustrates the relationship between the flanking identity score and the true benchmark sequence, using an example of a gap between two contigs of PAN027 on chromosome 2. The plot shows the edit distance (y-axis), which quantifies how different each candidate haplotype is from the benchmark. The red dot represents the true benchmark haplotype (PAN027.1), which achieves both a perfect flanking score and an edit distance of 0. The green dot indicates the result for T2T-CHM13, while the blue dot marks the haplotype selected by ImpuT2T. Overall, we observe a negative correlation between flanking score and edit distance, meaning that higher flanking scores generally correspond to haplotypes that are closer to the benchmark. Figure S14 **b** presents the flanking scores for the tie-break scenario, in which the flanking sequence length is extended to 25, 000 bp for the tied candidates identified in **a**. Although the haplotype with the highest flanking score does not always have the smallest edit distance to the benchmark, it is often a reasonable and effective choice.

#### 4.3.6 Filtering unlikely patching

After patch assembly, we applied a post-hoc quality filter at the chromosome level. Each patched scaffold combines donor contig sequence with inserted pangenome sequence spanning gaps between contigs. We evaluated three geometric and confidence-based criteria for patch sequence filtering.

Any connections with edge weight probability greater than 0.5 were protected from removal and the following filtering criteria are only for edges with weight lower than 0.5. First, the total length of inserted patch sequence must not exceed 10% of the combined length of the patched scaffold. Second, the longest single inserted patch sequence must not be longer than the shorter of the two flanking regions on either side of that insertion. Otherwise the patch sequence is removed. Third, any inserted patch longer than 2 Mb whose edge weight probability lower than 0.5 is treated as low-confidence and removed.

If any patch sequence has been removed, the scaffold is split that point. ImpuT2T recursively re-applied the same rules to the remaining parts. Fragments that still failed were dropped. Finally, isolated contigs not incorporated into a patched scaffold were retained only if their length was at least 10% of the shortest patched scaffold on that chromosome.

#### 4.3.7 Patching chromosome

In ImpuT2T, input contigs are provided as complete sequences. When patching a connection with a positive distance (i.e., a gap), the sequence bridging the gap is extracted from the selected donor haplotype and inserted between the two contigs. If the very ends of a contig are not represented in the alignment, a sequence of equal length is trimmed from the corresponding end of the patching haplotype to ensure a precise fit. For connections with negative distance (i.e., an overlap), the midpoint of the overlap is identified, and each contig is extended up to this midpoint so they can be merged seamlessly.

### 4.4 Experiment setup

Only the ImpuT2T patching workflows using the full panel, as well as the time and space subsampling experiments, were executed end-to-end with Snakemake. For all other subsampling experiments, we reused minimap alignment results from the full panel run, performing only the layout search, aggregation, and GQC evaluation steps.

#### 4.4.1 Superpopulation labels

Superpopulation labels for pangenome samples were assigned according to the International Genome Sample Resource (IGSR) [26]. For ten samples not covered by IGSR, we manually assigned superpopulation labels as follows: CN1 (EAS), YAO (EAS), HG002 (EUR), HG005 (EAS), T2T-CHM13 (EUR), GRCh38 (AFR), KSA001 (EUR), HG02109 (AFR), NA21309 (AFR), and HG06807 (AFR). We note that these superpopulation designations are used solely for analytical purposes and may not fully capture an individual’s ancestry.

#### 4.4.2 Benchmarking

The benchmark genomes for CN1, HG002, and HG005 were obtained from `A collection of high-quality human assemblies`. The PAN027 assembly was sourced from [16] (see Section 5). For HG005, we additionally scaffolded the HPRC2 assembly using YAO as a reference, using RagTag `scaffold` [13]. Any sequences that were not integrated into scaffolds were excluded from the benchmark set.

These benchmark genomes served not only as references for GQC evaluation, but also as data for training the logistic regression model. For model training and evaluation, the ImpuT2T workflow was run in a “single reference” pangenome, where the pangenome panel consisted solely of the corresponding benchmark genome, to generate a “groundtruth patch.” HG002’s groundtruth patch from the *de novo* assembly was used as the training set, while groundtruth patches from the other benchmark genomes were used as the test set.

Benchmarking of the patching on HG002 and PAN027 was performed using GQC with the default release resource configurations. For CN1 and HG005, benchmarking followed the GQC guidelines for new genomes, as described in the documentation under “Configuring GQC For New Genomes.”

For all four benchmark sets (HG002, PAN027, CN1, HG005), regions corresponding to the acrocentric short arms were excluded from the evaluation. The excluded region for each chromosome was defined as spanning from position 0 to the end of the Active Higher-Order Repeat (active_hor) plus 1 Mb.

For HG002, the default GQC configuration already includes an excluded BED file that marks regions of low-confidence, such as those composed of rDNA sequence arrays or potential misassemblies; we supplemented this by adding the acrocentric short arm regions to the configuration. Centromere satellite annotations for HG002 v1.1 were obtained from the Miga Lab CenSatData repository [27], generated using the standard T2T satellite profiling workflow [19].

For PAN027, centromere satellite annotations were taken from the official project repository [16, 28], and excluded regions were defined using the same approach, from chromosome start through active_hor plus 1 Mb.

For HG005, centromeric satellite annotations were sourced from the HPRC Freeze 2 data release [10]. Excluded regions were manually adjusted to align with the scaffold coordinates.

For CN1, we generated the acrocentric chromosome annotations using AniAnn’s [29], and excluded the short arms in a similar fashion.

See Section 5 for file paths.

## 5 Data availability

### 5.1 Reference assemblies

The HPRC2 assemblies utilized in this study were obtained from `A collection of high-quality human assemblies`, version 3.0 (https://zenodo.org/records/14854401). This collection comprises T2T-CHM13, GRCh38, CN1, YAO, KSA001, and 232 HPRC r2/v0.6 samples, resulting in a total of 472 haplotypes. Notably, this set forms a superset of the official HPRC release 2. For clarity and brevity, we refer to this entire collection as “HPRC2” throughout the manuscript.

Although HG002 and PAN027 (ID HG06807 in HPRC2) are also present in the HPRC2 dataset, we used the HG002 assembly from the Q100 project [30] and the PAN027 assembly from [16] as our benchmark references. For the HG005 benchmark, we scaffolded the HPRC2 HG005 assembly, with the corresponding AGP file provided in Supplementary Table S5. For CN1, the benchmark reference used is the same as that released as part of `A collection of high-quality human assemblies`.

### 5.2 Sequencing data

The donor HiFi sequencing data and parental short read for each sample were obtained from their respective sources as detailed below:

- **CN1**: https://download.cncb.ac.cn/gsa-human/HRA004405/ [6]
- **PAN027**: https://github.com/biomonika/HPP/tree/main/T2T-Pedigree-project%20 [16]
- **HG002**: https://s3-us-west-2.amazonaws.com/human-pangenomics/working/HPRC_PLUS/HG002/analysis/aligned_reads/hifi/GRCh38/HG002_aligned_GRCh38_winnowmap.sorted.bam [10]
- **HG005**: https://s3-us-west-2.amazonaws.com/human-pangenomics/working/HPRC_PLUS/HG005/analysis/aligned_reads/hifi/GRCh38/HG005_aligned_GRCh38_winnowmap.sorted.bam [10]

### 5.3 Annotation file

HG005 centromere annotation file is available at https://42basepairs.com/browse/s3/human-pangenomics/submissions/DC27718F-5F38-43B0-9A78-270F395F13E8--INT_ASM_PRODUCTION/HG005/assemblies/freeze_2/annotation/censat?file=HG005_pat_hprc_r2_v1.0.1.SatelliteStrand.bed (paternal) and https://42basepairs.com/browse/s3/human-pangenomics/submissions/DC27718F-5F38-43B0-9A78-270F395F13E8--INT_ASM_PRODUCTION/HG005/assemblies/freeze_2/annotation/censat?file=HG005_mat_hprc_r2_v1.0.1.SatelliteStrand.bed(maternal).

HG002 centromere annotation file is downloaded from the Miga Lab CenSatData repository https://github.com/hloucks/CenSatData

PAN027 centromere annotation file is downloaded from https://github.com/biomonika/washu-pedigree The excluded regions used for benchmarking CN1, as identified by AniAnn’s annotation, along with the excluded regions for all benchmark samples, are provided in Supplementary Tables S6–S9.

### 5.4 Sample list of the subsampled pangenome

The full lists of samples included in the balanced and population-specific subsampled pangenomes are provided in Supplementary Tables S10–S28. The “full 470” sample list are the full HPRC2 excluded the donor haplotypes.

## 6 Code availability

The program of ImpuT2T is available at https://github.com/maojanlin/ImpuT2T. The chromosome-partitioned HPRC2 is available through the download instruction in the same GitHub page or through Index zone https://lh3.github.io/s3bb/#bucket=https%3A%2F%2Fgenome-idx.s3.amazonaws.com&mode=folder&prefix=agc%2Fhprc2%2F.

## 7 Ethics approval

Not applicable.

## 10 Authors’ contributions

ML, VSS, and BL designed the method. ML wrote the software and performed the experiments. ML and BL wrote the manuscript. All authors read and approved the final manuscript.

## 11 Acknowledgments

This work was carried out at the Advanced Research Computing at Hopkins (ARCH) core facility, which is supported by the National Science Foundation (NSF) grant number OAC 1920103.

We thank Nancy Hansen for promptly answering questions and rapid updates to GQC, including releases supporting self-configuration for new genomes. We also thank Adam Phillippy for his valuable feedback and comments on this project.

We thank the AWS Open Data Sponsorship Program for sponsoring the “Index zone” S3 bucket (s3://genome-idx), where the chromosome-partitioned HPRC2 is hosted. We would like to acknowledge the National Human Genome Research Institute (NHGRI) for funding the following grants supporting the creation of the human pangenome reference: U41HG010972, U01HG010971, U01HG013760, U01HG013755, U01HG013748, U01HG013744, R01HG011274, and the Human Pangenome Reference Consortium (BioProject ID: PR-JNA730823)

## 12 Collaborators

*Human Pangenome Reference Consortium Version 2 Authors:* Derek Albracht^1^, Ivan A. Alexandrov^2^, Jamie Allen^3^, Alawi A. Alsheikh-Ali^4^, Nicolas Altemose^5^, Casey Andrews^6^, Dmitry Antipov^7^, Lucinda Antonacci-Fulton^1^, Mobin Asri^8^, Marcelo Ayllon^9^, Jennifer R. Balacco^10^, Floris P. Barthel^11^, Edward A. Belter Jr^1^, Halle D. Bender^8^, Andrew P. Blair^8^, Davide Bolognini^12^, Katherine E. Bonini^13^, Christina Boucher^14^, Guillaume Bourque^15,16,17^, Silvia Buonaiuto^18^, Shuo Cao^18^, Andrew Carroll^19^, Ann M. Mc Cartney^8^, Monika Cechova^8^, Mark J.P. Chaisson^20^, Pi-Chuan Chang^19^, Xian Chang^8^, Jitender Cheema^3^, Haoyu Cheng^21^, Claudio Ciofi^22^, Hiram Clawson^8^, Sarah Cody^1^, Vincenza Colonna^18^, Holland C. Conwell^23^, Robert Cook-Deegan^24^, Mark Diekhans^8^, Maria Angela Diroma^22^, Daniel Doerr^25,26,27^, Zheng Dong^6^, Danilo Dubocanin^5^, Richard Durbin^28,29^, Jana Ebler^25,30^, Evan E. Eichler^9,31^, Jordan M. Eizenga^8^, Parsa Eskandar^8^, Eddie Ferro^14^, Anna-Sophie Fiston-Lavier^32,33^, Sarah M. Ford^23^, Willard W. Ford^34^, Giulio Formenti^10^, Adam Frankish^3^, Mallory A. Freeberg^3^, Qichen Fu^6^, Stephanie M. Fullerton^35^, Robert S. Fulton^1^, Shenghan Gao^36^, Yan Gao^37^, Gage H. Garcia^9^, Obed A. Garcia^38^, Joshua M.V. Gardner^8^, Shilpa Garg^39^, Erik Garrison^18^, Nanibaa’ A. Garrison^40,41,42^, John E. Garza^1^, Margarita Geleta^43^, Mohammadmersad Ghorbani^44^, Tina A. Graves-Lindsay^1^, Richard E. Green^23^, Carol W. Greider^45^, Cristian Groza^46^, Bida Gu^20^, Andrea Guarracino^11,18^, Melissa Gymrek^47^, Maximilian Haeussler^8^, Leanne Haggerty^3^, Ira M. Hall^48,49^, Nancy F. Hansen^7^, Yue Hao^11^, Mohammad Amiruddin Hashmi^4^, David Haussler^8^, Prajna Hebbar^8^, Peter Heringer^25,26,27^, Glenn Hickey^8^, Todd L. Hillaker^8^, S. Nakib Hossain^3^, Neng Huang^37,50^, Sarah E. Hunt^3^, Toby Hunt^3^, Alexander G. Ioannidis^5,8^, Nafiseh Jafarzadeh^8^, Nivesh Jain^10^, Erich D. Jarvis^10,31^, Maryam Jehangir^11^, Juan Jiang^6^, Eimear E. Kenny^13^, Juhyun Kim^7^, Bonhwang Koo^10^, Sergey Koren^7^, Milinn Kremitzki^1,6^, Charles H. Langley^51^, Ben Langmead^52^, Heather A. Lawson^6^, Daofeng Li^6^, Heng Li^37,50^, Wen-Wei Liao^48,49^, Jiadong Lin^9^, Tianjie Liu^6^, Glennis A. Logsdon^36^, Ryan Lorig-Roach^8^, Jonathan LoTempio Jr^53^, Hailey Loucks^8^, Jane E. Loveland^3^, Jianguo Lu^54^, Shuangjia Lu^48,49^, Julian K. Lucas^8^, Walfred Ma^20^, Juan F. Macias-Velasco^1,6,55^, Kateryna D. Makova^56^, Maximillian G. Marin^37,50^, Christopher Markovic^1^, Tobias Marschall^25,30^, Franco L. Marsico^18^, Fergal J. Martin^3^, Mira Mastoras^8^, Capucine Mayoud^32^, Brandy McNulty^8^, Jack A. Medico^10^, Julian M. Menendez^8^, Karen H. Miga^8^, Anna Minkina^57^, Matthew W. Mitchell^58^, Saswat K. Mohanty^59^, Younes Mokrab^44,60,61^, Jean Monlong^62^, Shabir Moosa^44^, Avelina Moreno-Ochando^63,64^, Shinichi Morishita^65^, Jonathan M. Mudge^3^, Katherine M. Munson^9^, Njagi Mwaniki^66^, Nasna Nassir^4^, Chiara Natali^22^, Shloka Negi^8^, Lingbin Ni^9^, Adam M. Novak^8^, Faith Okamoto^8^, Keisuke K. Oshima^36^, Pilar N. Ossorio^67^, Chie Owa^65^, Sadye Paez^10^, Benedict Paten^8^, Clelia Peano^12,68^, Adam M. Phillippy^7^, Brandon D. Pickett^7^, Laura Pignata^18^, Nadia Pisanti^66^, David Porubsky^9,69^, Pjotr Prins^18^, Timofey Prodanov^25,30^, Anandi Radhakrishnan^8^, T. Rhyker Ranallo-Benavidez^11^, Brian J. Raney^8^, Mikko Rautiainen^70^, Alessandro Raveane^12^, Andreas Rechtsteiner^45^, Luyao Ren^9,31^, Arang Rhie^7^, Fedor Ryabov^71,72^, Samuel Sacco^23^, Farnaz Salehi^18^, Michael C. Schatz^52,73^, Laura B. Scheinfeldt^74^, Aarushi Sehgal^34^, William E. Seligmann^23^, Mahsa Shabani^75^, Kishwar Shafin^19^, Shadi Shahatit^32^, Ruhollah Shemirani^13^, Vikram S. Shivakumar^52^, Swati Sinha^3^, Jouni Sirén^8^, Linnéa Smeds^59^, Steven J. Solar^7^, Marco Sollitto^10,22^, Nicole Soranzo^12,28,76^, Andrew B. Stergachis^9,57^, Marie-Marthe Suner^3^, Yoshihiko Suzuki^65^, Arda Sö ylev^25,30^, Ahmad Abou Tayoun^77,78^, Jack A.S. Tierney^3^, Chad Tomlinson^1^, Francesca Floriana Tricomi^3^, Mohammed Uddin^4,79^, Matteo Tommaso Ungaro^23,80^, Rahul Varki^14^, Flavia Villani^18^, Ivo Violich^8^, Mitchell R. Vollger^57^, Brian P. Walenz^7^, Charles Wang^81^, Lisa E. Wang^13^, Ting Wang^1,6,55^, Aaron M. Wenger^82^, Conor V. Whelan^10^, Zilan Xin^6^, Zheng Xu^6^, Kai Ye^83^, DongAhn Yoo^9^, Wenjin Zhang^6^, Ying Zhou^37^, Xiaoyu Zhuo^6^, Giulia Zunino^12^

*Affiliations:*

^1^McDonnell Genome Institute, Washington University School of Medicine, St. Louis, MO 63108, USA

^2^Department of Human Molecular Genetics and Biochemistry, Faculty of Medical and Health Sciences, Tel Aviv University, Tel Aviv 69978, Israel

^3^European Molecular Biology Laboratory, European Bioinformatics Institute (EMBL-EBI), Wellcome Genome Campus, Hinxton, Cambridge CB10 1SD, UK

^4^Center for Applied and Translational Genomics (CATG), Mohammed Bin Rashid University of Medicine and Health Sciences, Dubai Health, Dubai, UAE

^5^Department of Genetics, Stanford University, Palo Alto, CA 94304 USA

^6^Department of Genetics, Washington University School of Medicine, St. Louis, MO 63110, USA

^7^Genome Informatics Section, Center for Genomics and Data Science Research, National Human Genome Research Institute, National Institutes of Health, Bethesda, MD 20892, USA

^8^UC Santa Cruz Genomics Institute, University of California, Santa Cruz, CA 95060, USA

^9^Department of Genome Sciences, University of Washington School of Medicine, Seattle, WA 98195, USA

^10^The Vertebrate Genome Laboratory, The Rockefeller University, New York, NY 10065, USA

^11^Bioinnovation and Genome Sciences, The Translational Genomics Research Institute (TGen), Phoenix, AZ 85004, USA

^12^Human Technopole, Milan, Italy

^13^Institute for Genomic Health, Icahn School of Medicine at Mount Sinai, New York, NY 10029, USA

^14^Department of Computer and Information Science and Engineering, University of Florida, Gainesville, FL 32611, USA

^15^Canadian Center for Computational Genomics, McGill University, Montréal, QC H3A 0G1, Canada

^16^Department of Human Genetics, McGill University, Montréal, QC H3A 0G1, Canada

^17^Victor Phillip Dahdaleh Institute of Genomic Medicine, Montréal, QC H3A 0G1, Canada

^18^Department of Genetics, Genomics and Informatics, University of Tennessee Health Science Center, Memphis, TN 38163, USA

^19^Google LLC, Mountain View, CA 94043, USA

^20^Quantitative and Computational Biology, University of Southern California, Los Angeles, CA 90089, USA

^21^Department of Biomedical Informatics and Data Science, Yale School of Medicine, New Haven, CT 06510, USA

^22^Department of Biology, University of Florence, Sesto Fiorentino, FI 50019, Italy

^23^Department of Ecology and Evolutionary Biology, University of California, Santa Cruz, CA 95060, USA

^24^Arizona State University, Consortium for Science, Policy & Outcomes, Washington, DC 20006, USA

^25^Center for Digital Medicine, Heinrich Heine University Dü sseldorf, Dü sseldorf, NRW, DE

^26^Department for Endocrinology and Diabetology at the Medical Faculty and University Hospital Dü sseldorf, Heinrich Heine University Dü sseldorf, Dü sseldorf, NRW, DE

^27^Paul-Langerhans-Group Computational Diabetology, German Diabetes Center (DDZ) and Leibniz Institute for Diabetes Research, Dü sseldorf, NRW, DE

^28^Wellcome Sanger Institute, Genome Campus, Hinxton, CB10 1RQ, UK

^29^Department of Genetics, University of Cambridge, Cambridge, CB2 3EH, UK

^30^Institute for Medical Biometry and Bioinformatics, Medical Faculty and University Hospital Dü sseldorf, Heinrich Heine University, Dü sseldorf, NRW, DE

^31^Howard Hughes Medical Institute, Chevy Chase, MD 20815, USA

^32^ISEM, Univ Montpellier, CNRS, IRD, Montpellier, FR

^33^Institut Universitaire de France, Paris, FR

^34^Department of Computer Science and Engineering, University of California San Diego, La Jolla, CA 92093, USA

^35^Department of Bioethics & Humanities, University of Washington School of Medicine, Seattle, WA 98195, USA

^36^Department of Genetics, Epigenetics Institute, Perelman School of Medicine, University of Pennsylvania, Philadelphia, PA 19104, USA

^37^Department of Data Science, Dana-Farber Cancer Institute, Boston, MA 02215, USA

^38^Department of Anthropology, University of Kansas, Lawrence, KS 66045, USA

^39^School of Health Sciences, University of Manchester, Manchester M13 9PL, UK

^40^Traditional, ancestral and unceded territory of the Gabrielino/Tongva peoples, Institute for Society & Genetics, University of California, Los Angeles, Los Angeles, CA 90095, USA

^41^Traditional, ancestral and unceded territory of the Gabrielino/Tongva peoples, Institute for Precision Health, David Geffen School of edicine, University of California, Los Angeles, Los Angeles, CA 90095, USA

^42^Traditional, ancestral and unceded territory of the Gabrielino/Tongva peoples, Division of General Internal Medicine & Health Services Research, David Geffen School of Medicine, University of California, Los Angeles, Los Angeles, CA 90095, USA

^43^Department of Electrical Engineering and Computer Science, University of California, Berkeley, Berkeley, CA 94720, USA

^44^Medical and Population Genomics Lab, Sidra Medicine, Doha, Qatar

^45^Department of Molecular Cell and Developmental Biology, University of California, Santa Cruz, CA, USA

^46^Montreal Heart Institute, Montréal, QC, Canada

^47^Department of Pediatrics, University of California San Diego, La Jolla, CA 92093, USA

^48^Center for Genomic Health, Yale University School of Medicine, New Haven, CT 06510, USA

^49^Department of Genetics, Yale University School of Medicine, New Haven, CT 06510, USA

^50^Department of Biomedical Informatics, Harvard Medical School, Boston, MA 02115, USA

^51^Department of Evolution and Ecology and the Center for Population Biology, University of California, One Shields, Davis, CA 95616, USA

^52^Department of Computer Science, Johns Hopkins University, Baltimore, MD 21218, USA

^53^Department of Pediatrics, Division of Genetics, School of Medicine, University of California, Irvine, CA 92697, USA

^54^Sun Yat-sen University, Guangzhou, China

^55^Edison Family Center for Genome Sciences & Systems Biology, Washington University School of Medicine, St. Louis, MO 63110, USA

^56^Department of Biology and Center for Medical Genomics, Penn State University, University Park, PA 16802, USA

^57^Division of Medical Genetics, Department of Medicine, University of Washington School of Medicine, Seattle, WA 98195, USA

^58^The Jackson Laboratory for Genomic Medicine, Farmington, CT 06032, USA

^59^Department of Biology, Penn State University, University Park, PA 16802, USA

^60^Department of Biomedical Science, College of Health Sciences, Qatar University, Doha, Qatar

^61^Department of Genetic Medicine, Weill Cornell Medicine-Qatar, Doha, Qatar

^62^IRSD - Digestive Health Research Institute, University of Toulouse, INSERM, INRAE, ENVT, UPS, Toulouse, FR

^63^MATCH biosystems, S.L., Elche, Spain

^64^Universidad Miguel Hernández de Elche, Elche, Spain

^65^Department of Computational Biology and Medical Sciences, The University of Tokyo, Kashiwa, Chiba 277-8561, Japan

^66^Department of Computer Science, University of Pisa, Pisa, Italy

^67^Law School, University of Wisconsin-Madison, Madison, WI 53706, USA

^68^Institute of Genetics and Biomedical Research, UoS of Milan, National Research Council, Milan, Italy

^69^Genome Biology Unit, European Molecular Biology Laboratory (EMBL), Heidelberg, DE

^70^Institute for Molecular Medicine Finland, Helsinki Institute of Life Science, University of Helsinki, Helsinki, Finland

^71^The Center for Bio- and Medical Technologies, Moscow, RUS

^72^Centre for Biomedical Research and Technology, HSE University, Moscow, RUS

^73^Department of Biology, Johns Hopkins University, Baltimore, MD 21218, USA

^74^Coriell Institute for Medical Research, Camden, NJ 08103, USA

^75^University of Amsterdam, Amsterdam, Netherlands

^76^School of Clinical Medicine, University of Cambridge, Cambridge, CB2 0SP, UK

^77^Center for Genomic Discovery, Mohammed Bin Rashid University, Dubai Health, UAE

^78^Dubai Health Genomic Medicine Center, Dubai Health, UAE

^79^GenomeArc Inc, Mississauga, ON, Canada

^80^Department of Biology and Biotechnologies “Charles Darwin”, University of Rome “La Sapienza”, Rome 00185, IT

^81^Center for Genomics, Loma Linda University School of Medicine, Loma Linda, CA 92350, USA

^82^PacBio, Menlo Park, CA 94025, USA

^83^The first affiliated hospital of Xi’an Jiaotong University, Xi’an Jiaotong University, Xi’an, Shaanxi, 710049, China

## References

[1] Jacob Pritt, Nae-Chyun Chen, and Ben Langmead. FORGe: prioritizing variants for graph genomes. Genome Biology, 19(1):220, 2018.

[2] Mao-Jan Lin, Sheila Iyer, Nae-Chyun Chen, and Ben Langmead. Measuring, visualizing, and diagnosing reference bias with biastools. Genome Biology, 25(1):101, 2024.

[3] Haoyu Cheng, Gregory T Concepcion, Xiaowen Feng, Haowen Zhang, and Heng Li. Haplotype-resolved de novo assembly using phased assembly graphs with hifiasm. Nature Methods, 18(2):170–175, 2021.

[4] S. Nurk, S. Koren, A. Rhie, M. Rautiainen, A. V. Bzikadze, A. Mikheenko, M. R. Vollger, N. Altemose, L. Uralsky, A. Gershman, S. Aganezov, S. J. Hoyt, M. Diekhans, G. A. Logsdon, M. Alonge, S. E. Antonarakis, M. Borchers, G. G. Bouffard, S. Y. Brooks, G. V. Caldas, N. C. Chen, H. Cheng, C. S. Chin, W. Chow, L. G. de Lima, P. C. Dishuck, R. Durbin, T. Dvorkina, I. T. Fiddes, G. Formenti, R. S. Fulton, A. Fungtam-masan, E. Garrison, P. G. S. Grady, T. A. Graves-Lindsay, I. M. Hall, N. F. Hansen, G. A. Hartley, M. Haukness, K. Howe, M. W. Hunkapiller, C. Jain, M. Jain, E. D. Jarvis, P. Kerpedjiev, M. Kirsche, M. Kolmogorov, J. Korlach, M. Kremitzki, H. Li, V. V. Maduro, T. Marschall, A. M. McCartney, J. McDaniel, D. E. Miller, J. C. Mullikin, E. W. Myers, N. D. Olson, B. Paten, P. Peluso, P. A. Pevzner, D. Porubsky, T. Potapova, E. I. Rogaev, J. A. Rosenfeld, S. L. Salzberg, V. A. Schneider, F. J. Sedlazeck, K. Shafin, C. J. Shew, A. Shumate, Y. Sims, A. F. A. Smit, D. C. Soto, I. ć, J. M. Storer, A. Streets, B. A. Sullivan, F. Thibaud-Nissen, J. Torrance, J. Wagner, B. P. Walenz, A. Wenger, J. M. D. Wood, C. Xiao, S. M. Yan, A. C. Young, S. Zarate, U. Surti, R. C. McCoy, M. Y. Dennis, I. A. Alexandrov, J. L. Gerton, R. J. O’Neill, W. Timp, J. M. Zook, M. C. Schatz, E. E. Eichler, K. H. Miga, and A. M. Phillippy. The complete sequence of a human genome. Science, 376(6588):44–53, Apr 2022.

[5] Nancy F Hansen, Nathan Dwarshuis, Hyun Joo Ji, Arang Rhie, Hailey Loucks, Glennis A Logsdon, Mitchell R Vollger, Jessica M Storer, Juhyun Kim, Eleni Adam, et al. A complete diploid human genome benchmark for personalized genomics. bioRxiv, 2025.

[6] Chentao Yang, Yang Zhou, Yanni Song, Dongya Wu, Yan Zeng, Lei Nie, Panhong Liu, Shilong Zhang, Guangji Chen, Jinjin Xu, et al. The complete and fully-phased diploid genome of a male Han Chinese. Cell Research, 33(10):745–761, 2023.

[7] Yukun He, Yanan Chu, Shuming Guo, Jiang Hu, Ran Li, Yali Zheng, Xinqian Ma, Zhenglin Du, Lili Zhao, Wenyi Yu, et al. T2T-YAO: a telomere-to-telomere assembled diploid reference genome for Han Chinese. Genomics, Proteomics & Bioinformatics, 21 (6):1085–1100, 2023.

[8] Maxat Kulmanov, Rund Tawfiq, Yang Liu, Hatoon Al Ali, Marwa Abdelhakim, Mohammed Alarawi, Hind Aldakhil, Dana Alhattab, Ebtehal A Alsolme, Azza Althagafi, et al. A reference quality, fully annotated diploid genome from a Saudi individual. Scientific Data, 11(1):1278, 2024.

[9] Prasad Sarashetti, Josipa Lipovac, Qi Jia, Ling Wang, Zhe Li, Lovro Vrcek, Ying Chen, Fei Yao, YY Sia, Dehui Lin, et al. A complete Telomere-to-Telomere diploid reference genome for South Asian population. bioRxiv, pages 2025–07, 2025.

[10] Wen-Wei Liao, Mobin Asri, Jana Ebler, Daniel Doerr, Marina Haukness, Glenn Hickey, Shuangjia Lu, Julian K Lucas, Jean Monlong, Haley J Abel, et al. A draft human pangenome reference. Nature, 617(7960):312–324, 2023.

[11] Glennis A Logsdon, Peter Ebert, Peter A Audano, Mark Loftus, David Porubsky, Jana Ebler, Feyza Yilmaz, Pille Hallast, Timofey Prodanov, DongAhn Yoo, et al. Complex genetic variation in nearly complete human genomes. Nature, 644(8076):430–441, 2025.

[12] Yifei Wang, Zhongqu Duan, Dan Chen, Dandan Shi, Yi Ding, Zhibin Wang, Bao-qing Li, Zhiyi Wang, Minmin Guo, Wen Yang, et al. The 1000 Chinese Pangenome empowers medical and population genetics. Nature, pages 1–10, 2026.

[13] Michael Alonge, Ludivine Lebeigle, Melanie Kirsche, Katie Jenike, Shujun Ou, Sergey Aganezov, Xingang Wang, Zachary B Lippman, Michael C Schatz, and Sebastian Soyk. Automated assembly scaffolding using RagTag elevates a new tomato system for high-throughput genome editing. Genome Biology, 23(1):258, 2022.

[14] Adam Diehl and Alan Boyle. Fast and accurate draft genome patching with GPatch. bioRxiv, 2025.

[15] Julian K Lucas, Prajna Hebbar, Wen-Wei Liao, Juan F Macias-Velasco, Adam M Novak, Mobin Asri, Jennifer R Balacco, Andrew P Blair, Jana Ebler, Joshua MV Gardner, et al. Hprc2: A human pangenome reference with near-complete coverage of common genetic variation. bioRxiv, pages 2026–07, 2026.

[16] Monika Cechova, Tamara A Potapova, Andreas Rechtsteiner, Glenn Hickey, Rebecca Serra Mari, Mira Mastoras, Julian Menendez, Nikol Poláková, Prajna Hebbar, Fedor Ryabov, et al. Complete genomes of a multi-generational pedigree to expand studies of genetic and epigenetic inheritance. bioRxiv, pages 2025–12, 2025.

[17] Heng Li. Minimap2: pairwise alignment for nucleotide sequences. Bioinformatics, 34 (18):3094–3100, 2018.

[18] Heng Li. auN: a new metric to measure assembly contiguity. Heng Li’s blog, April

[19] Nicolas Altemose, Glennis A Logsdon, Andrey V Bzikadze, Pragya Sidhwani, Sasha A Langley, Gina V Caldas, Savannah J Hoyt, Lev Uralsky, Fedor D Ryabov, Colin J Shew, et al. Complete genomic and epigenetic maps of human centromeres. Science, 376(6588):eabl4178, 2022.

[20] Steven J Solar, Prajna Hebbar, Leonardo Gomes de Lima, Alex Sweeten, Arang Rhie, Tamara Potapova, Luciana de Gennaro, Andrea Guarracino, Juhyun Kim, Brandon D Pickett, et al. Origin and evolution of acrocentric chromosomes in human and great apes. bioRxiv, pages 2025–12, 2025.

[21] Rahul Varki, Massimiliano Rossi, Eddie Ferro, Marco Oliva, Erik Garrison, Ben Langmead, and Christina Boucher. Accurate short-read alignment through r-index-based pangenome indexing. Genome Research, 35(7):1609, 2025.

[22] Xian Chang, Adam M Novak, Jordan M Eizenga, Jouni Sirén, Jean Monlong, Shloka Negi, Francesco Andreace, Sagorika Nag, Konstantinos Kyriakidis, Glenn Hickey, et al. Rapid, accurate long-and short-read mapping to large pangenome graphs with vg Giraffe. bioRxiv, pages 2025–09, 2025.

23. Jouni Sirén, Parsa Eskandar, Matteo Tommaso Ungaro, Glenn Hickey, Jordan M Eizenga, Adam M Novak, Xian Chang, Pi-Chuan Chang, Mikhail Kolmogorov, Andrew Carroll, et al. Personalized pangenome references. Nature Methods, 21(11): 2017–2023, 2024.

[24] Glenn Hickey, Jean Monlong, Jana Ebler, Adam M Novak, Jordan M Eizenga, Yan Gao, Tobias Marschall, Heng Li, and Benedict Paten. Pangenome graph construction from genome alignments with minigraph-cactus. Nature biotechnology, 42(4):663–673, 2024.

[25] Vikram S Shivakumar and Ben Langmead. Mumemto: efficient maximal matching across pangenomes. Genome Biology, 26(1):169, 2025.

[26] Susan Fairley, Ernesto Lowy-Gallego, Emily Perry, and Paul Flicek. The international genome sample resource (igsr) collection of open human genomic variation resources. Nucleic acids research, 48(D1):D941–D947, 2020.

27. Hailey Loucks and Miga Lab. CenSatData: Centromere and satellite annotations for genomic benchmarks. https://github.com/hloucks/CenSatData, 2026.

[28] Monika Cechova and Miga Lab. washu-pedigree: Complete genomes of a multi-generational pedigree to expand studies of genetic and epigenetic inheritance. https://github.com/biomonika/washu-pedigree, 2026.

[29] Alexander P Sweeten, Michael Schatz, and Adam M Phillippy. Aniann’s: alignment-free annotation of tandem repeat arrays using fast average nucleotide identity estimates. bioRxiv, pages 2026–01, 2026.

[30] Mikko Rautiainen, Sergey Nurk, Brian P Walenz, Glennis A Logsdon, David Porubsky, Arang Rhie, Evan E Eichler, Adam M Phillippy, and Sergey Koren. Telomere-to-telomere assembly of diploid chromosomes with Verkko. Nature Biotechnology, pages 1–9, 2023.

