## Supplementary Figures S1-S14 and Supplementary Table S1-S4 for "ImpuT2T: Pangenome-Based Patching for Human Genome Assemblies"

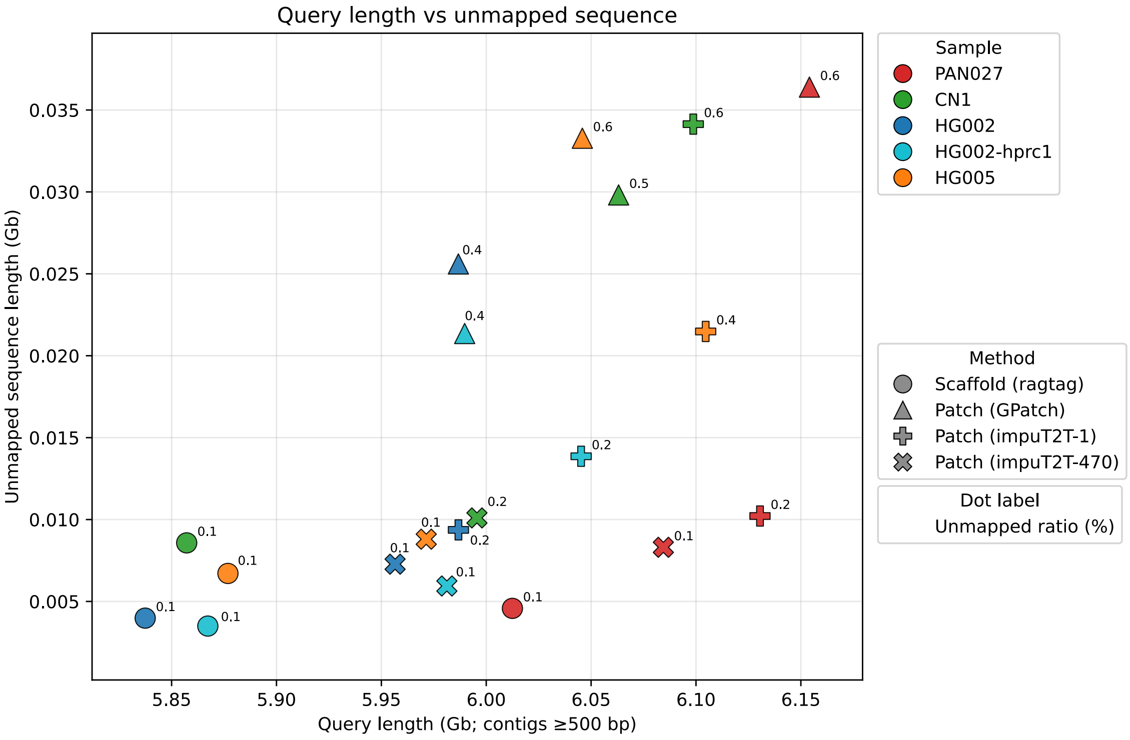


Figure S1: Query sequence length vs the total unmapped query sequence for each of the five assemblies with different enhancing methods. The label next to each dot shows the unmapped ratio of each query genome.

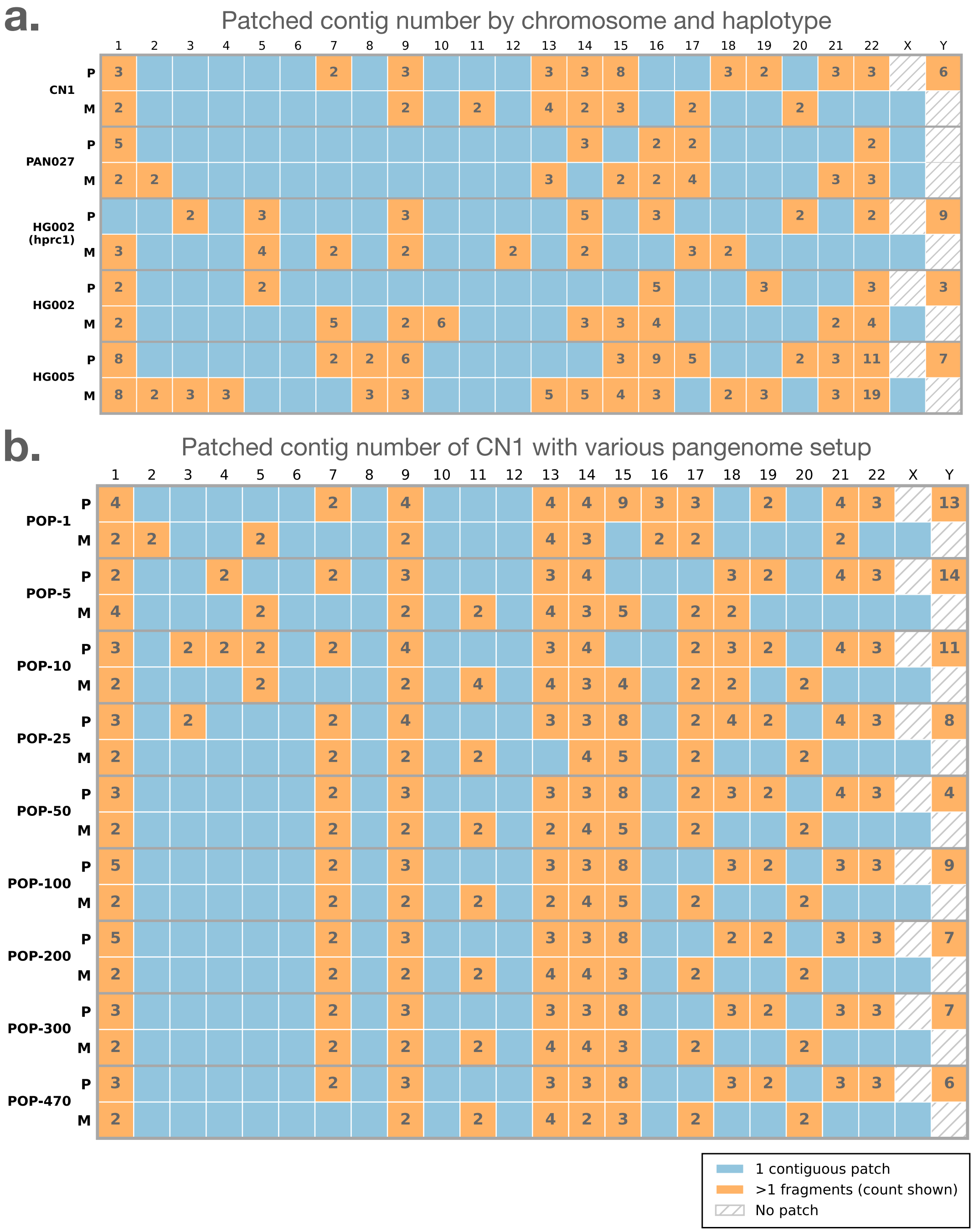


Figure S2: **a**. Number of chromosome/haplotype fragments in each of the five test datasets after patching with ImpuT2T using the full HPRC panel. **b**. Number of chromosome/haplotype fragments for CN1 after patching with ImpuT2T using varying subset of the HPRC2 pangenome.

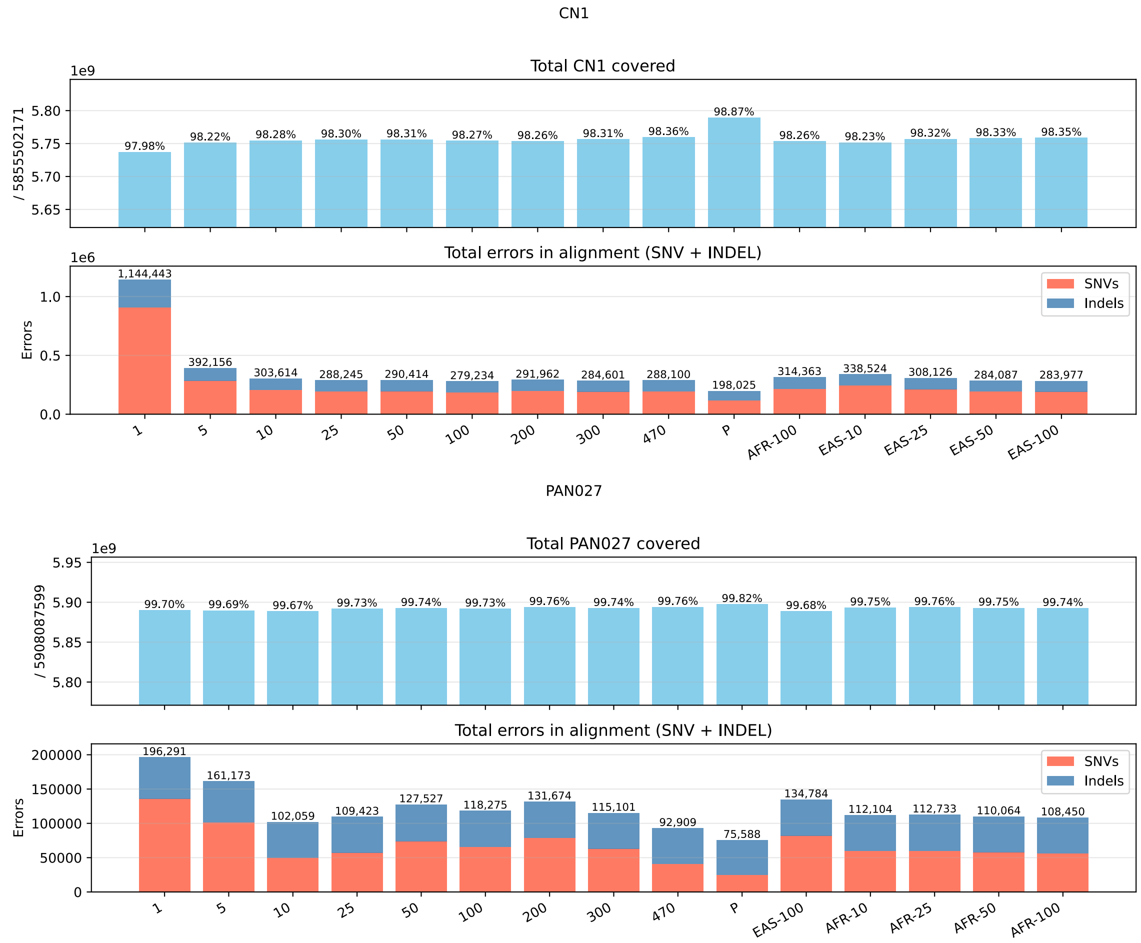


Figure S3: Summary of benchmark coverage and error counts for the CN1 and PAN027 assemblies. **Top:** Bar plot displaying coverage statistics and error counts for CN1. **Bottom:** Bar plot displaying coverage statistics and error counts for PAN027.

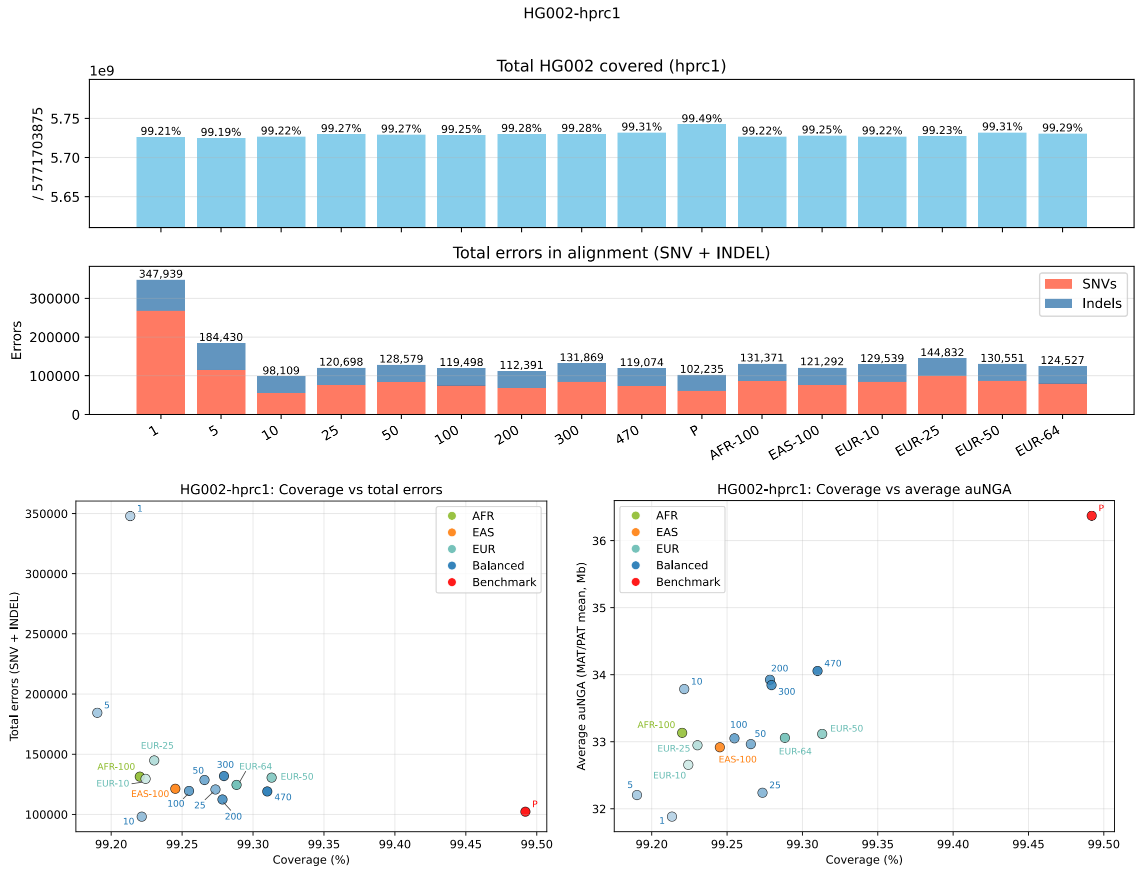


Figure S4: Overview of benchmark results for HPRC1 HG002 patched with HPRC2 subsets. EUR is the matching population to HG002. **Top:** Bar plot summarizing coverage statistics and error counts across various pangenome patching strategies for HG002-HPRC1. **Bottom Left:** Scatter plot showing the relationship between coverage and total error counts for each patching strategies. **Bottom Right:** Scatter plot displaying the relationship between coverage and average auNGA across each patching strategies.

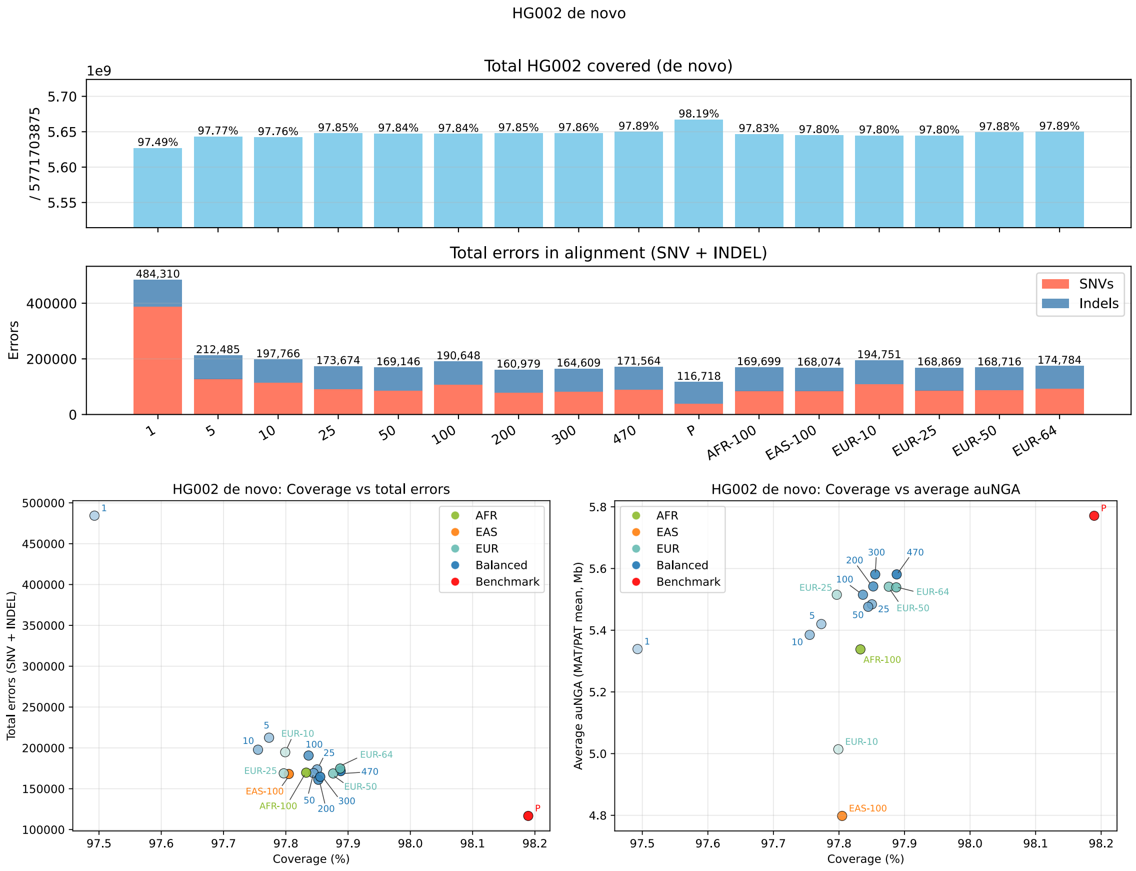


Figure S5: Overview of benchmark results for *de novo* assembled HG002 patched with HPRC2 subsets. EUR is the matching population to HG002. **Top:** Bar plot summarizing coverage statistics and error counts across various pangenome patching strategies for HG002. **Bottom Left:** Scatter plot showing the relationship between coverage and total error counts for each patching strategies. **Bottom Right:** Scatter plot displaying the relationship between coverage and average auNGA across each patching strategies.

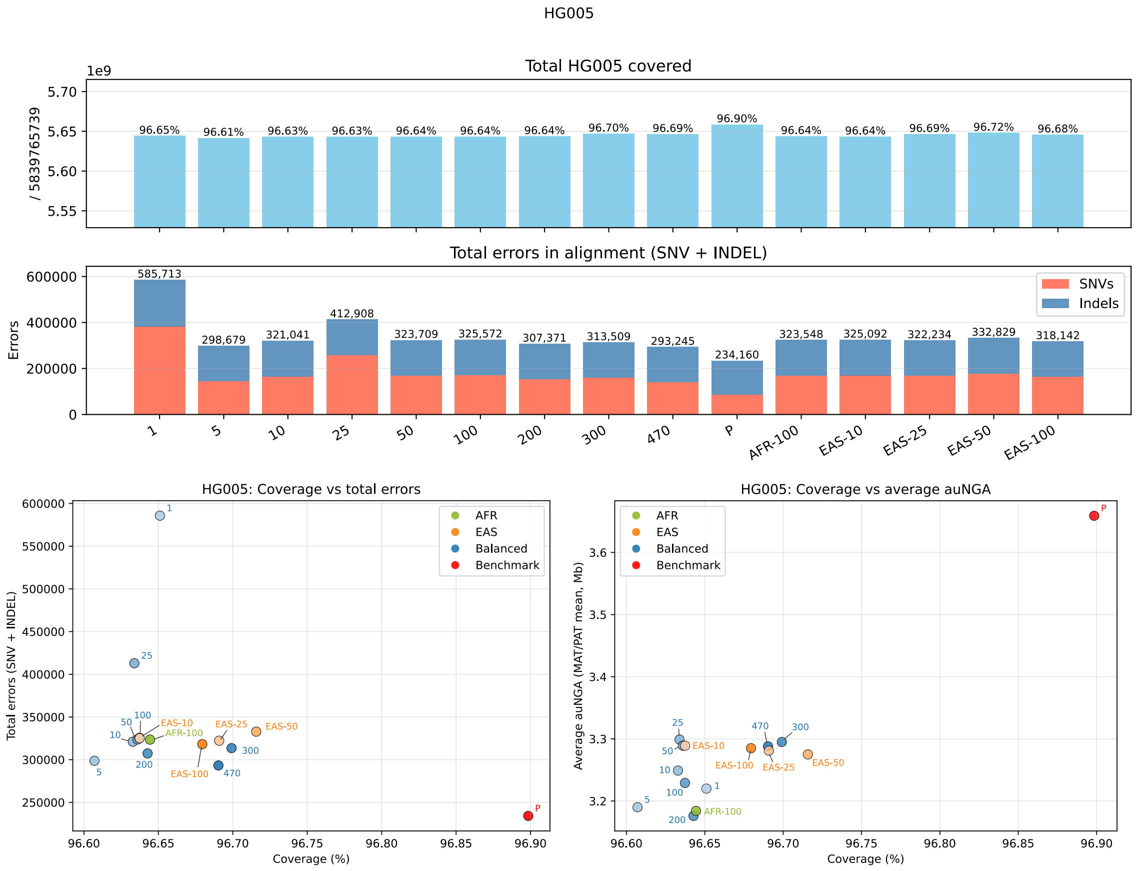


Figure S6: Overview of benchmark results for HG005 patched with HPRC2 subsets. EAS is the matching population to HG005. **Top:** Bar plot summarizing coverage statistics and error counts across various pangenome patching strategies for HG005. **Bottom Left:** Scatter plot showing the relationship between coverage and total error counts for each patching strategies. **Bottom Right:** Scatter plot displaying the relationship between coverage and average auNGA across each patching strategies.

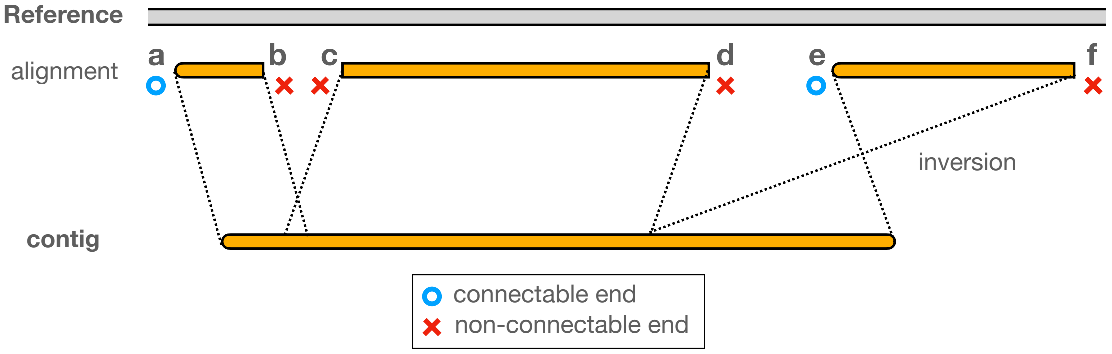


Figure S7: A contig can be split into several aligned segments when mapped to the reference. Only those segment ends that overlap the terminal regions of a contig are considered "connectable" and eligible for assembly connections. In this illustration, connectable ends are marked with blue circles, while non-connectable ends are indicated by red crosses. Of note, even though ends **a** and **e** are connectable, they can only be joined to sequence on their left relative to the alignment on reference genome.

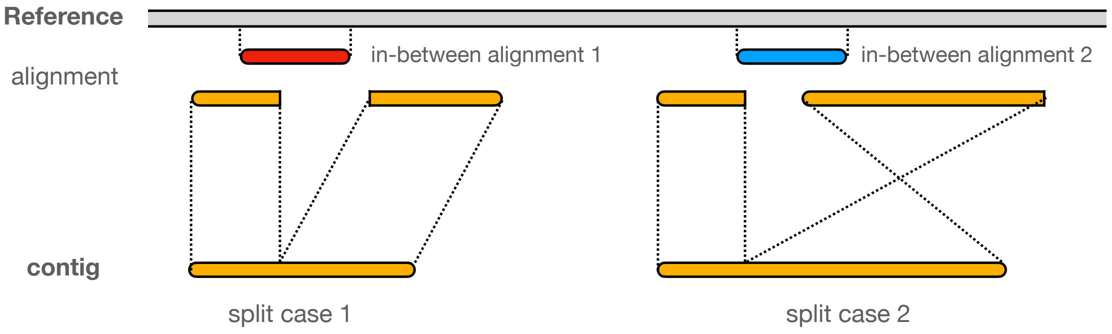


Figure S8: Illustration of how split contig alignments are interpreted based on segment orientation. In **split case 1**, both aligned segments are in the same orientation and therefore considered to represent a single contiguous alignment. As a result, the **in-between alignment 1** (red) is "contained" within this combined mapping. In contrast, in **split case 2**, the two aligned segments are in opposite orientations and are treated as two separate segments; consequently, the **in-between alignment 2** (blue) is not "contained" in this scenario.

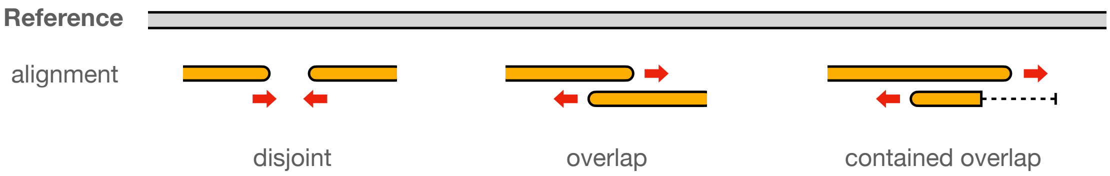


Figure S9: A connection between two contig segments can be established when their respective connectable ends are oriented toward each other. These connectable segment ends can be classified into three categories: disjoint, overlapping, or contained overlap. In the contained overlap scenario, the "contained" segment must be part of a contig whose full length extends beyond the outer boundaries of the overlapping segment.

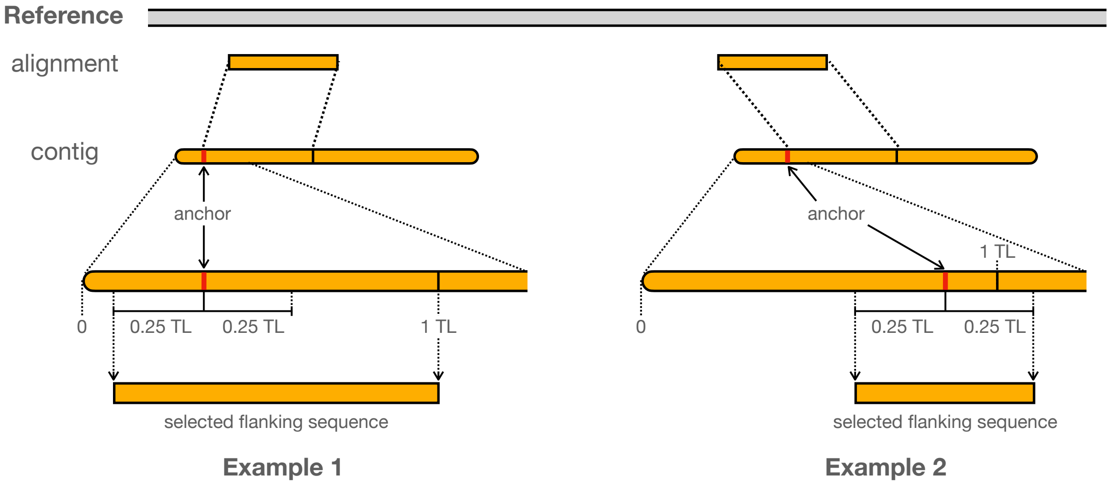


Figure S10: Schematic illustrating the selection of flanking sequence for terminal alignment. The anchor point is defined as the end of the aligned segment nearest to the contig edge. The terminal length (TL) is set to $5,000$ bp by default, or extended to $25,000$ bp for tie-breaks. A symmetric window of $\pm\frac{1}{4}$ TL is centered on the anchor within the contig. For the left flanking sequence, the selected region spans from $\max(0, \text{anchor} - \frac{1}{4}\text{TL})$ to $\max(\text{anchor} + \frac{1}{4}\text{TL}, \text{TL})$. The right flanking sequence is defined analogously, mirroring this procedure at the right end of the contig.

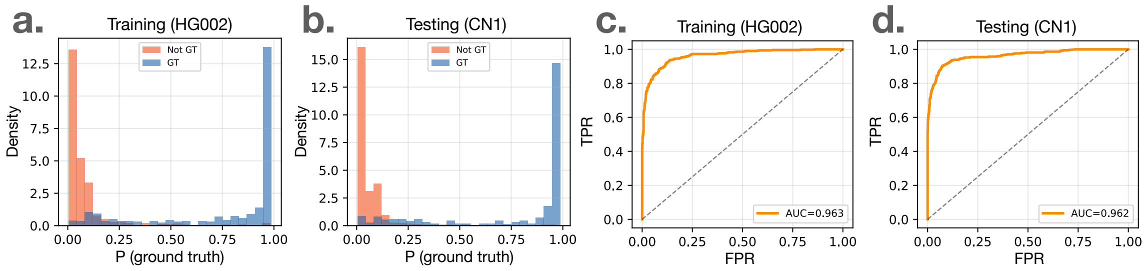


Figure S11: Overview of ImpuT2T's internal logistic regression model. **a** and **b**: Histograms of prediction score distributions for edges that do or do not belong to the ground truth, shown for both the training set (HG002) and an independent test set (CN1). **e** and **f**: Relationship between flanking sequence similarity scores and edit distance to the benchmark sequence, using default flanking sequence length of $5,000$ bp (**e**) and extended length of $25,000$ bp (**f**).

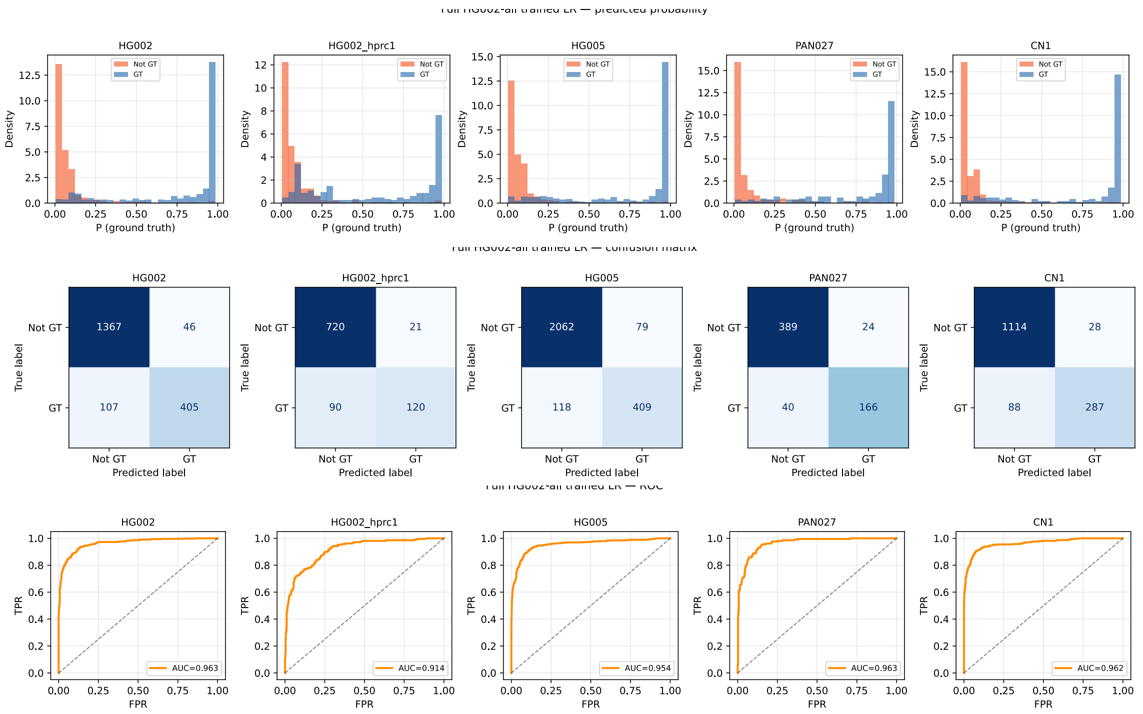


Figure S12: From top to bottom are the probability histograms, confusion matrices, and the ROC curves of the logistic regression model on the five test dataset, *de novo* assembled HG002, HG002 from HPRC1, HG005, PAN027, and CN1. The model was trained on *de novo* assembled HG002.

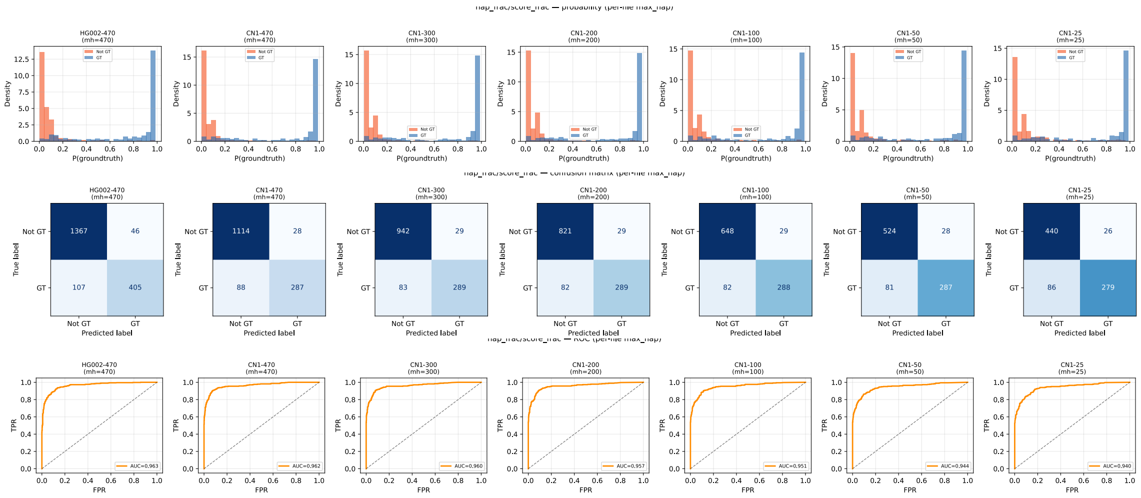


Figure S13: From top to bottom are the probability histograms, confusion matrices, and the ROC curves of the logistic regression model on the training *de novo* assembled HG002, and CN1 test dataset with different size of the pangenome. The size of the pangenome for CN1 are arranged from $470$, $300$, $200$, $100$, $50$, to $25$.

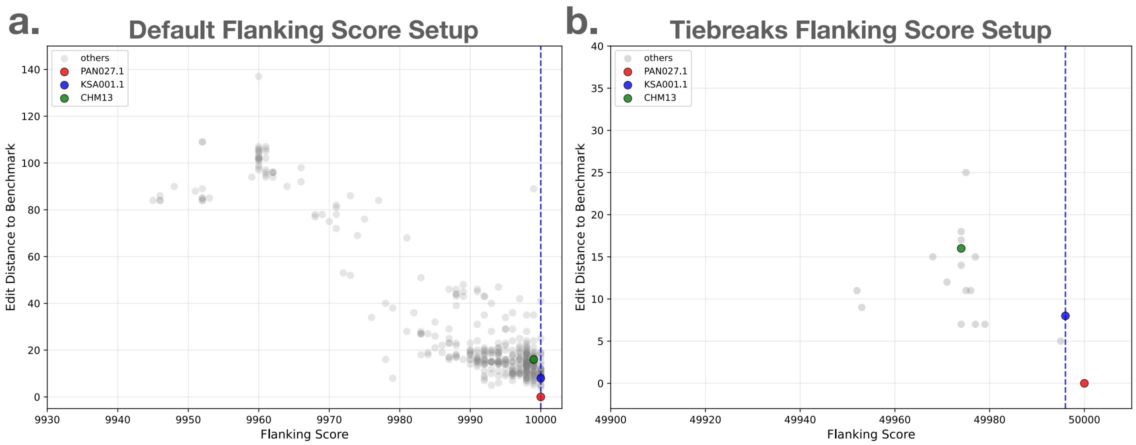


Figure S14: Relationship between flanking sequence similarity scores and edit distance to the benchmark sequence, using default flanking sequence length of $5,000$ bp (**a**) and extended length of $25,000$ bp (**b**).

PAN027.1: the benchmark sample in this patched gap.

### Supplementary Tables

Table S1: Comparison capitategories of chromosomes across the five dataset

| | CN1 | PAN027 | HG002-HPRC1 | HG002 | HG005 | Total |
|---|---|---|---|---|---|---|
| **Err & Cov better** | 10 | 4 | 6 | 5 | 5 | 30 (13.2%) |
| **Err better only** | 12 | 13 | 18 | 15 | 18 | 76 (33.3%) |
| **Cov better only** | 6 | 3 | 2 | 0 | 5 | 16 (7.0%) |
| **Trade-off** | 4 | 1 | 2 | 2 | 2 | 11 (4.8%) |
| **Comparable** | 12 | 19 | 13 | 21 | 12 | 77 (33.8%) |
| **Err worse only** | 1 | 2 | 4 | 2 | 3 | 12 (5.3%) |
| **Cov worse only** | 1 | 4 | 0 | 0 | 0 | 5 (2.2%) |
| **Err & Cov worse** | 0 | 0 | 0 | 0 | 1 | 1 (0.4%) |

Table S2: Per chromosome-haplotype error counts (indels / SNVs) across methods and assemblies. For each entry, the lower value between ImpuT2T and GPatch is indicated in bold.

| Chr- | CN1 | | | PAN027 | | | HG002-HPRC1 | | | HG002 | | | HG005 | | |
|---|---|---|---|---|---|---|---|---|---|---|---|---|---|---|---|
| Hap | scaff | ImpuT2T | GPatch | scaff | ImpuT2T | GPatch | scaff | ImpuT2T | GPatch | scaff | ImpuT2T | GPatch | scaff | ImpuT2T | GPatch |
| 1-P | 3988/9587 | 5073/**26107** | **4975**/29337 | 2032/503 | **2001/500** | 2161/3578 | 1604/877 | **1659/1130** | 1746/5749 | 3174/986 | **3263/1158** | 5661/69328 | 5964/1515 | **6200/1446** | 7054/32067 |
| 1-M | 6810/26503 | **6992/28544** | 8406/64378 | 2206/2381 | **2183/1773** | 3344/38462 | 1698/1303 | **1752/1633** | 3238/60832 | 3259/839 | **3670/2374** | 6831/106116 | 6301/2220 | **6285/1614** | 8290/53203 |
| 2-P | 2961/2482 | **3046/2687** | 3350/6008 | 2041/1954 | **2082/2773** | 2537/6337 | 1782/1377 | **1849/1495** | 2694/22301 | 3318/2530 | **3507/3032** | 3966/20234 | 5364/732 | **5447/880** | 5757/5304 |
| 2-M | 2710/893 | **2770/929** | 3263/4526 | 1654/115 | **1829/492** | 3082/29650 | 1700/494 | **1735/591** | 3526/35701 | 3262/1459 | **3419/1780** | 4031/23285 | 5942/3052 | **5999/2846** | 6137/4805 |
| 3-P | 2271/2598 | **2702/8695** | 4070/40707 | 1449/780 | **1466/893** | 2276/21291 | 1215/85 | **1215/85** | 2230/20384 | 2371/1774 | **2670/5169** | 3119/16526 | 4466/4745 | **4522/5144** | 6079/31394 |
| 3-M | 2279/1134 | **2303/1725** | 3353/28581 | 1459/233 | **1459/197** | 3155/43525 | 1352/1418 | **1369/1747** | 2721/30354 | 2313/306 | **2459/2600** | 2977/13200 | 5040/12046 | **5130/12669** | 7188/72525 |
| 4-P | 2023/2709 | **2327/4002** | 3158/10355 | 1966/10844 | **1980/11704** | 2535/24495 | 1142/75 | **1606/2995** | 1719/14499 | 2492/3269 | 2608/**3775** | **2567**/11268 | 3918/2681 | **3970/3612** | 5794/43011 |
| 4-M | 2205/8337 | **2418/9377** | 2683/13269 | 1528/2330 | 1878/4684 | **1548/2602** | 1184/161 | **1224/539** | 1766/12947 | 2173/1034 | **2167/1180** | 2877/12604 | 3846/1415 | **3973**/4320 | 4040/**2950** |
| 5-P | 2069/1455 | 2265/2241 | **2224/2011** | 1424/613 | 1466/**621** | **1447**/1097 | 1074/727 | 1366/**4596** | **1304**/4650 | 2072/461 | **2295/2397** | 2608/13507 | 3997/1970 | **4096/1248** | 4153/2611 |
| 5-M | 2151/2857 | 2444/6114 | **2335/3664** | 1289/227 | 1303/**234** | **1294**/357 | 1065/1126 | 1131/1838 | **1108/1280** | 2047/248 | **2057/440** | 2130/738 | 4044/648 | **4044/861** | 4133/975 |
| 6-P | 1835/573 | **1850/574** | 2041/1967 | 1200/1364 | 1214/1605 | **1173/395** | 1088/1526 | **1104/2034** | 1836/51550 | 2021/2494 | **2230/7393** | 2837/48915 | 3722/1077 | 3764/**873** | **3756**/1087 |
| 6-M | 1860/1355 | **1866/1247** | 2093/7650 | 1239/1118 | **1260/217** | 1325/8151 | 1093/1693 | 1193/5194 | **1158/2387** | 2181/2680 | 2203/**3263** | **2174**/3299 | 3833/616 | **3876/496** | 4380/9699 |
| 7-P | 2067/1119 | **2592/2610** | 2896/19365 | 1500/131 | **1507/143** | 1832/991 | 1360/5229 | **1267/4546** | 1415/6298 | 2087/447 | **2208/508** | 2349/913 | 4079/1796 | **4160/1671** | 4204/1981 |
| 7-M | 2183/988 | **2952/3454** | 3404/14327 | 1295/101 | **1330/165** | 1351/283 | 1124/141 | **1124/155** | 1149/617 | 2538/1205 | **2601/1266** | 2623/1497 | 4225/799 | **4260/693** | 4471/1500 |
| 8-P | 1739/2138 | **1971/5013** | 2068/5331 | 1069/74 | **1254**/2414 | 1333/**1578** | 926/68 | **1084/1912** | 1265/4447 | 1848/1624 | **1779/665** | 1970/2368 | 3334/2514 | **3314/1159** | 3600/4526 |
| 8-M | 1597/962 | **1686/1277** | 1860/2820 | 1219/1002 | **1254/1309** | 1555/3675 | 965/256 | **1095/1089** | 1180/2894 | 1894/1081 | **1890/1058** | 1932/1739 | 3424/1013 | **3432/730** | 3803/4377 |
| 9-P | 1657/1992 | **2151/3564** | 24068/74482 | 935/74 | **936/74** | 63794/216203 | 956/511 | **1661/1540** | 3262/5590 | 1836/976 | **1789/1493** | 5637/7680 | 3218/591 | **3394/689** | 34128/58254 |
| 9-M | 2007/3644 | **2362/4653** | 21035/66296 | 1023/985 | **1041/1020** | 12543/38157 | 1647/3400 | **3771/8409** | 18614/48452 | 2680/2195 | **2849/8240** | 34507/77972 | 3132/629 | **3229/1751** | 5374/15120 |
| 10-P | 1876/3197 | 1902/**3216** | **1872**/3224 | 1104/149 | **1118**/421 | 1173/**286** | 856/97 | 841/191 | **825/92** | 1671/363 | **1744**/1199 | 1746/**398** | 3332/760 | **3328/785** | 3439/1069 |
| 10-M | 1645/1038 | 1700/1151 | **1681/1046** | 1021/78 | **1024/160** | 1042/228 | 865/186 | **925/265** | 1026/1231 | 1689/208 | 1819/799 | **1764/323** | 3334/359 | 3482/960 | **3347/362** |
| 11-P | 1556/374 | 1837/1705 | **1718/941** | 990/83 | 998/117 | **994/88** | 908/81 | **924/453** | 948/3911 | 1606/241 | **1652/1895** | 1660/4498 | 3210/420 | **3292/764** | 3583/23194 |
| 11-M | 1658/2257 | 2021/3968 | **1968/3466** | 899/320 | **903/532** | 1258/21625 | 936/570 | **936/570** | 1505/34744 | 1743/328 | **1781/1784** | 2388/36072 | 3234/1087 | **3237/1533** | 4667/88699 |
| 12-P | 1771/1289 | **1839/1422** | 1883/1719 | 1077/66 | **1087/68** | 1112/443 | 918/289 | **918/111** | **918**/289 | 1780/302 | **1805/310** | 1808/378 | 3448/412 | 3459/419 | **3442/413** |
| 12-M | 1710/727 | 1779/1008 | **1767/917** | 1245/225 | 1245/**225** | **1204**/277 | 889/417 | **889/417** | **889/417** | 1853/217 | **1855/219** | 1896/309 | 3538/396 | 3617/2594 | **3562/401** |
| 13-P | 981/189 | 995/**189** | **986**/193 | 690/74 | 703/77 | **693/75** | 585/53 | **595/63** | 617/138 | 1097/126 | **1105/139** | **1105**/178 | 2033/262 | 2034/**264** | **2031**/462 |
| 13-M | 1001/141 | **1001**/142 | 1002/**141** | 684/59 | 686/60 | **684/59** | 634/685 | 805/2261 | **599/311** | 1294/1629 | **1269/1259** | 1322/1898 | 2103/260 | **2102/260** | 2110/283 |
| 14-P | 1515/2753 | **1596/3197** | 1600/3401 | 751/127 | **748/88** | 783/448 | 646/444 | 930/2603 | **673/705** | 1344/1707 | **1480/2486** | 1504/2893 | 2154/294 | **2153/312** | 2214/983 |
| 14-M | 1226/863 | 1377/1447 | **1269/965** | 911/129 | 951/**131** | **908**/351 | 1086/4145 | **834/2182** | 1135/4461 | 1142/473 | 1416/2432 | **1102/172** | 2313/464 | **2393/280** | 2403/533 |
| 15-P | 1517/2629 | 1597/3028 | **1543/2781** | 726/56 | **726/56** | **726/56** | 585/41 | 586/**41** | **585/41** | 1188/421 | 1203/451 | **1186/444** | 2427/1636 | 3033/4782 | **2691/2556** |
| 15-M | 1330/2703 | 1845/9527 | **1350/2688** | 783/72 | **783/72** | **783/72** | 588/63 | **588/62** | 591/72 | 1217/1236 | 1220/1187 | **1170/266** | 3893/15320 | **5619/27789** | 14189/117713 |
| 16-P | 1135/2330 | **1424/2957** | 2656/27632 | 885/192 | **950/702** | 3009/58167 | 1049/1691 | **1582/7770** | 2349/40341 | 1566/1515 | **1648/1040** | 2677/30792 | 2736/278 | **3227**/7820 | 3262/**5909** |
| 16-M | 1305/3169 | **1734/4457** | 3268/42103 | 1049/456 | **1156**/3638 | 1227/**2594** | 762/582 | **771/473** | 1219/6298 | 1567/288 | **1916/2226** | 1923/2899 | 2962/2049 | **3840/12414** | 5471/59164 |
| 17-P | 1767/1592 | **2171/2595** | 6666/74844 | 989/64 | **995/64** | 1090/735 | 981/3904 | 1122/**4861** | **988**/4886 | 1848/4417 | **1783/3263** | 1922/5359 | 2834/322 | **3046/3674** | 3290/10473 |
| 17-M | 1559/987 | **2116/2529** | 2946/16864 | 951/932 | **938/95** | 1506/9452 | 784/201 | **803/773** | 1073/11251 | 1531/591 | **1638/2612** | 1704/5311 | 3236/3211 | **3400**/7341 | 3608/**7250** |
| 18-P | 1092/4801 | **1062**/4037 | 1098/**3628** | 674/68 | **693/93** | 729/182 | 746/3711 | **738/3438** | 1201/10121 | 938/566 | **1187/2715** | 1661/9712 | 1874/2644 | **1891/2816** | 1967/3004 |
| 18-M | 968/2059 | **924/2265** | 1273/5540 | 542/76 | **557/80** | 649/2920 | 633/2065 | **590/1255** | 639/2600 | 989/400 | **996/473** | 1051/1373 | 1803/1417 | **1869/2495** | 2068/4383 |
| 19-P | 1332/854 | **3176/4657** | 3192/5049 | 976/64 | 1017/76 | **972/69** | 590/2394 | 604/2361 | **587/403** | 1533/324 | **1540/331** | 1621/3443 | 2687/1302 | **2716/1371** | 2767/1627 |
| 19-M | 1344/1055 | **3006/4137** | 3872/6363 | 1186/634 | **1201/662** | 1273/1021 | 587/77 | **612/52** | 888/10968 | 1545/121 | **1614/4193** | 2107/19587 | 2840/571 | **3215**/6057 | 3237/**2213** |
| 20-P | 1075/927 | **1146/1096** | 1200/1292 | 677/147 | **728/153** | 743/799 | 829/7615 | **450/652** | 817/7319 | 1291/7811 | 1292/7809 | **1279/7648** | 1537/749 | **1530/304** | 1630/1957 |
| 20-M | 1249/6620 | **1033/792** | 1339/6856 | 585/74 | **590/104** | 612/213 | 374/31 | **375/31** | 376/32 | 915/144 | 990/1544 | **924/214** | 1754/1269 | **1824/939** | 2157/11733 |
| 21-P | 477/137 | 561/342 | **508/281** | 227/10 | **227/10** | **227/10** | 214/17 | **214/17** | **214/17** | 453/53 | **455/53** | 464/65 | 792/86 | **829**/98 | 836/**90** |
| 21-M | 411/1499 | **424**/1605 | 460/**216** | 253/11 | **253/11** | **253/11** | 251/23 | **251/23** | **251/23** | 453/55 | 482/**67** | **457/67** | 813/105 | 825/**104** | **816**/124 |
| 22-P | 648/326 | **823/794** | 861/926 | 442/435 | **448/438** | **448**/446 | 244/17 | **244/17** | **244/17** | 621/92 | **625/94** | 635/106 | 1148/90 | **1158/93** | 1159/103 |
| 22-M | 661/182 | **996/1166** | 1110/1665 | 420/186 | 425/**184** | **417/184** | 247/11 | 248/**15** | **247/15** | 666/100 | **667/103** | **667**/105 | 1208/103 | 1235/166 | **1207/101** |
| X-P | – | – | – | 1628/212 | **1723/439** | 2776/3930 | – | – | – | – | – | – | – | – | – |
| X-M | 1571/592 | **2267/9934** | 3073/18335 | 1706/930 | **1882/1137** | 2999/5759 | 581/163 | **678/330** | 1146/1853 | 1638/365 | **1766/478** | 2148/2330 | 3582/507 | **3684/1889** | 4070/2638 |
| Y-P | 719/3794 | **1740/8062** | 9863/143722 | – | – | – | – | – | – | – | – | – | 684/3700 | **1750/9333** | 12373/187597 |

Table S3: Coverage by chromosome-haplotype across methods and assemblies. The maximum value for each entry is shown in bold.

| Chr-Hap | CN1 | | | PAN027 | | | HG002-HPRC1 | | | HG002 | | | HG005 | | |
|---|---|---|---|---|---|---|---|---|---|---|---|---|---|---|---|
| | **scaff** | **ImpuT2T** | **GPatch** | **scaff** | **ImpuT2T** | **GPatch** | **scaff** | **ImpuT2T** | **GPatch** | **scaff** | **ImpuT2T** | **GPatch** | **scaff** | **ImpuT2T** | **GPatch** |
| 1-P | 0.9658 | **0.9771** | 0.9657 | **0.9995** | 0.9995 | 0.9995 | 0.9853 | **0.9953** | 0.9853 | 0.9589 | **0.9782** | 0.9422 | 0.9611 | **0.9653** | 0.9489 |
| 1-M | 0.9648 | **0.9668** | 0.9616 | 0.9884 | **0.9953** | 0.9892 | 0.9909 | **0.9979** | 0.9840 | 0.9831 | **0.9848** | 0.9846 | 0.9596 | **0.9650** | 0.9555 |
| 2-P | 0.9939 | **0.9950** | 0.9876 | 0.9896 | **0.9940** | 0.9872 | 0.9990 | **0.9996** | 0.9987 | 0.9770 | **0.9774** | 0.9765 | 0.9583 | **0.9598** | 0.9596 |
| 2-M | 0.9927 | **0.9936** | 0.9936 | 0.9977 | **0.9991** | 0.9988 | 0.9966 | **0.9973** | 0.9927 | 0.9864 | **0.9883** | 0.9830 | 0.9756 | 0.9770 | **0.9780** |
| 3-P | 0.9899 | **0.9935** | 0.9933 | 0.9969 | **0.9990** | 0.9833 | 0.9870 | 0.9869 | **0.9905** | 0.9702 | **0.9726** | 0.9725 | 0.9530 | **0.9565** | 0.9553 |
| 3-M | 0.9936 | 0.9943 | **0.9955** | **0.9987** | 0.9985 | 0.9987 | 0.9929 | **0.9959** | 0.9935 | 0.9797 | **0.9829** | 0.9796 | 0.9777 | **0.9780** | 0.9772 |
| 4-P | 0.9813 | 0.9862 | **0.9873** | 0.9962 | 0.9962 | **0.9963** | **0.9991** | **0.9991** | 0.9990 | 0.9851 | **0.9874** | 0.9857 | 0.9686 | **0.9703** | 0.9700 |
| 4-M | 0.9905 | 0.9922 | **0.9930** | 0.9972 | 0.9937 | **0.9973** | 0.9968 | **0.9973** | 0.9972 | 0.9888 | 0.9880 | **0.9897** | 0.9688 | **0.9701** | 0.9654 |
| 5-P | 0.9891 | 0.9920 | **0.9922** | 0.9993 | 0.9993 | **0.9993** | 0.9570 | **0.9787** | 0.9589 | 0.9346 | **0.9526** | 0.9393 | 0.9558 | **0.9617** | 0.9583 |
| 5-M | 0.9917 | **0.9934** | 0.9925 | 0.9993 | **0.9993** | 0.9993 | 0.9926 | **0.9931** | 0.9929 | **0.9843** | 0.9842 | 0.9806 | 0.9718 | **0.9731** | 0.9693 |
| 6-P | 0.9972 | **0.9974** | 0.9972 | 0.9988 | **0.9990** | 0.9973 | 0.9772 | 0.9781 | **0.9794** | 0.9573 | **0.9726** | 0.9592 | **0.9657** | 0.9649 | 0.9657 |
| 6-M | 0.9936 | 0.9938 | **0.9962** | 0.9985 | 0.9997 | **0.9997** | 0.9735 | **0.9776** | 0.9739 | 0.9656 | **0.9664** | 0.9659 | 0.9584 | 0.9601 | **0.9640** |
| 7-P | 0.9746 | **0.9807** | 0.9713 | 0.9989 | **0.9991** | 0.9989 | 0.9899 | 0.9924 | **0.9937** | 0.9774 | **0.9792** | 0.9782 | **0.9638** | 0.9635 | 0.9632 |
| 7-M | 0.9799 | 0.9873 | **0.9878** | 0.9978 | **0.9992** | 0.9979 | 0.9977 | **0.9979** | 0.9978 | 0.9858 | **0.9877** | 0.9869 | 0.9798 | 0.9788 | **0.9803** |
| 8-P | 0.9939 | 0.9966 | **0.9975** | 0.9916 | **0.9982** | 0.9916 | 0.9864 | **0.9955** | 0.9954 | 0.9794 | 0.9766 | **0.9794** | 0.9623 | **0.9668** | 0.9666 |
| 8-M | 0.9945 | 0.9954 | **0.9958** | **0.9969** | 0.9939 | 0.9935 | 0.9869 | **0.9945** | 0.9890 | 0.9812 | 0.9810 | **0.9812** | 0.9775 | 0.9735 | **0.9776** |
| 9-P | 0.9512 | **0.9741** | 0.9602 | 0.9988 | 0.9988 | **0.9990** | **0.9816** | 0.9797 | 0.9798 | 0.9746 | **0.9797** | 0.9772 | 0.9627 | **0.9708** | 0.9283 |
| 9-M | 0.9634 | **0.9800** | 0.9570 | 0.9981 | **0.9989** | 0.9979 | **0.9890** | 0.9874 | 0.9884 | 0.9794 | **0.9838** | 0.9810 | 0.9668 | **0.9670** | 0.9639 |
| 10-P | 0.9969 | **0.9976** | 0.9976 | 0.9983 | 0.9983 | **0.9983** | 0.9966 | 0.9965 | **0.9966** | 0.9674 | 0.9750 | **0.9754** | **0.9620** | 0.9605 | 0.9618 |
| 10-M | 0.9978 | **0.9983** | 0.9909 | 0.9991 | **0.9991** | 0.9991 | 0.9902 | **0.9978** | 0.9962 | 0.9795 | **0.9850** | 0.9803 | 0.9749 | 0.9739 | **0.9778** |
| 11-P | 0.9746 | **0.9810** | 0.9779 | 0.9988 | **0.9988** | 0.9988 | 0.9977 | **0.9986** | 0.9978 | 0.9738 | **0.9758** | 0.9743 | 0.9661 | 0.9668 | **0.9676** |
| 11-M | 0.9790 | **0.9896** | 0.9872 | 0.9944 | **0.9994** | 0.9982 | 0.9980 | 0.9980 | **0.9983** | 0.9810 | **0.9815** | 0.9811 | 0.9690 | **0.9719** | 0.9574 |
| 12-P | 0.9863 | **0.9894** | 0.9893 | 0.9994 | **0.9994** | 0.9994 | 0.9963 | **0.9963** | 0.9963 | 0.9767 | **0.9774** | 0.9771 | 0.9657 | **0.9663** | 0.9615 |
| 12-M | 0.9827 | **0.9827** | 0.9827 | **0.9992** | **0.9992** | 0.9992 | **0.9960** | 0.9960 | **0.9960** | 0.9881 | **0.9882** | 0.9881 | 0.9752 | 0.9752 | **0.9753** |
| 13-P | 0.9138 | 0.9137 | **0.9139** | 0.8997 | **0.8998** | 0.8997 | 0.9966 | 0.9966 | **0.9980** | 0.9843 | **0.9846** | 0.9845 | 0.8933 | **0.8935** | 0.8935 |
| 13-M | **0.9161** | **0.9161** | **0.9161** | 0.8795 | **0.8795** | 0.8795 | 0.9969 | **0.9976** | 0.9973 | 0.9885 | 0.9884 | **0.9887** | 0.9141 | **0.9146** | 0.9146 |
| 14-P | 0.8860 | **0.8928** | 0.8875 | **0.8753** | 0.8752 | 0.8752 | 0.9891 | 0.9908 | **0.9952** | 0.9789 | **0.9800** | 0.9788 | 0.7843 | **0.7846** | 0.7845 |
| 14-M | 0.8944 | **0.9030** | 0.8997 | 0.8882 | **0.8882** | 0.8882 | 0.9799 | **0.9965** | 0.9799 | 0.9773 | 0.9776 | **0.9783** | 0.9498 | 0.9499 | **0.9499** |
| 15-P | 0.8826 | **0.8915** | 0.8830 | **0.8312** | **0.8312** | **0.8312** | 0.9908 | **0.9908** | 0.9908 | 0.9762 | 0.9755 | **0.9767** | 0.8306 | **0.8337** | 0.8313 |
| 15-M | 0.8334 | **0.8424** | 0.8333 | **0.8381** | **0.8381** | **0.8381** | 0.9960 | 0.9960 | **0.9960** | 0.9861 | **0.9876** | 0.9870 | 0.9440 | **0.9577** | 0.9526 |
| 16-P | 0.9575 | 0.9629 | **0.9652** | 0.9944 | 0.9961 | **0.9987** | 0.9717 | **0.9927** | 0.9850 | 0.9445 | **0.9726** | 0.9718 | 0.9585 | 0.9583 | **0.9653** |
| 16-M | 0.9491 | **0.9740** | 0.9641 | **0.9980** | 0.9974 | 0.9968 | **0.9960** | 0.9943 | 0.9690 | 0.9804 | 0.9807 | **0.9826** | 0.9608 | 0.9727 | **0.9749** |
| 17-P | 0.9274 | **0.9457** | 0.9438 | 0.9978 | **0.9993** | 0.9978 | 0.9700 | **0.9911** | 0.9750 | 0.9547 | **0.9677** | 0.9645 | 0.9294 | 0.9390 | **0.9533** |
| 17-M | 0.9232 | **0.9606** | 0.9553 | 0.9794 | 0.9795 | **0.9957** | 0.9840 | 0.9852 | **0.9907** | 0.9475 | 0.9536 | **0.9591** | 0.9254 | **0.9582** | 0.9450 |
| 18-P | 0.9852 | **0.9870** | 0.9847 | 0.9977 | **0.9986** | 0.9978 | 0.9873 | 0.9867 | **0.9950** | 0.9730 | **0.9773** | 0.9754 | 0.9721 | **0.9741** | 0.9735 |
| 18-M | 0.9724 | 0.9757 | **0.9836** | 0.9973 | 0.9978 | **0.9979** | 0.9926 | 0.9915 | **0.9950** | 0.9801 | 0.9804 | **0.9832** | 0.9757 | **0.9798** | 0.9798 |
| 19-P | 0.8467 | 0.9020 | **0.9026** | 0.9998 | **0.9999** | 0.9712 | 0.9912 | **0.9920** | 0.9917 | 0.9503 | **0.9610** | 0.9488 | 0.9400 | 0.9399 | **0.9402** |
| 19-M | 0.8480 | 0.8984 | **0.9092** | 0.9984 | 0.9985 | **0.9995** | 0.9974 | 0.9970 | **0.9975** | 0.9748 | 0.9710 | **0.9777** | 0.9462 | 0.9487 | **0.9496** |
| 20-P | 0.9764 | 0.9846 | **0.9847** | 0.9993 | 0.9992 | **0.9993** | 0.9877 | **0.9893** | 0.9881 | **0.9524** | 0.9521 | 0.9493 | **0.9420** | 0.9400 | 0.9390 |
| 20-M | 0.9819 | **0.9854** | 0.9836 | 0.9979 | 0.9974 | **0.9980** | **0.9844** | 0.9844 | 0.9843 | **0.9858** | 0.9848 | 0.9858 | 0.9694 | **0.9721** | 0.9696 |
| 21-P | 0.7839 | **0.7891** | 0.7883 | **0.7833** | **0.7833** | **0.7833** | **0.9966** | **0.9966** | **0.9966** | **0.9788** | **0.9788** | 0.9788 | 0.7892 | **0.7894** | 0.7893 |
| 21-M | 0.7549 | 0.7552 | **0.7557** | **0.7658** | **0.7658** | **0.7658** | **1.0000** | **1.0000** | **1.0000** | **0.9917** | 0.9913 | 0.9890 | **0.7487** | 0.7486 | 0.7486 |
| 22-P | 0.6664 | **0.6667** | 0.6647 | 0.7687 | **0.7687** | 0.7687 | **0.9987** | 0.9987 | **0.9987** | 0.9721 | **0.9722** | **0.9722** | 0.7181 | 0.7185 | **0.7185** |
| 22-M | 0.6372 | **0.6614** | 0.6606 | **0.7500** | **0.7500** | **0.7500** | 0.9930 | **0.9948** | **0.9948** | 0.9536 | 0.9510 | **0.9538** | **0.6982** | 0.6978 | 0.6973 |
| X-P | – | – | – | 0.9966 | **0.9972** | 0.9934 | – | – | – | – | – | – | – | – | – |
| X-M | 0.9620 | **0.9811** | 0.9750 | 0.9935 | **0.9952** | 0.9898 | 0.9986 | **0.9999** | 0.9958 | 0.9962 | **0.9984** | 0.9946 | 0.9949 | **0.9970** | 0.9922 |
| Y-P | 0.7442 | **0.9142** | 0.8206 | – | – | – | – | – | – | – | – | – | 0.7231 | 0.8771 | **0.8810** |

Table S4: Sequencing statistics for the donor HiFi and parental short reads used in hifiasm trio-binning *de novo* assemblies. Coverage is total bases divided by an assumed haploid genome size of 3.1 Gbp.

| **Sample** | **HiFi bases (Gbp)** | **HiFi N50 (bp)** | **HiFi cov. (×)** | **Maternal cov. (×)** | **Paternal cov. (×)** |
|---|---|---|---|---|---|
| **HG002** | 110.55 | 14,718 | 35.7 | 32.7 | 32.8 |
| **HG005** | 145.34 | 16,956 | 46.9 | 55.8 | 54.1 |
| **CN1** | 182.31 | 14,149 | 58.8 | 57.2 | 46.6 |
| **PAN027** | 120.87 | 19,670 | 39.0 | 37.8 | 40.3 |
